# Computational design of a thermostable α-helical barrel protein with tunable active-site positioning

**DOI:** 10.1101/2024.10.07.617055

**Authors:** Wael Elaily, David Stoll, Morakot Chakatok, Matteo Aleotti, Birgit Grill, Horst Lechner, Mélanie Hall, Gustav Oberdorfer

## Abstract

Over the last decades, transformative catalytic strategies have emerged, with biocatalysis currently exerting a substantial influence on the pharmaceutical and fine chemical sectors. Fast progress in the design of efficient enzymatic processes, however, suffers from the lack of readily available, stable, and customizable protein scaffolds that can be adapted to different catalytic functions. Here, we detail the design and experimental characterization of computationally designed de novo proteins with a non-natural fold for biocatalytic applications. The initial design and several variants form a helical barrel structure comprised of six antiparallel straight helices connected by five loops, creating an open central channel with two accessible cavities. To demonstrate the versatility of this scaffold, we designed variants with catalytic sites positioned at different locations along the central channel. All designs show high thermal stability and excellent agreement between experimental and calculated scattering profiles from small-angle X-ray scattering, while a crystal structure of a surface-redesigned variant confirms the close match between the designed and experimental structures. Importantly, repositioning and engineering the catalytic sites enables substantial modulation of catalytic activity, with the best variant showing an approximately 11-fold increase in catalytic efficiency compared with the original design. Finally, the designs can be used for whole-cell biotransformations and tolerate up to 20% organic solvent. These results establish a stable de novo protein scaffold with tunable functional sites, offering a versatile platform for biocatalysis, biosensing, and biosynthetic systems.

## Introduction

Since the turn of the century, biocatalysis has emerged as an indispensable tool for chemists, offering unparalleled precision and selectivity in chemical transformations^1^. Particularly in the production of fine chemicals, biocatalysis has become a method of choice, owing to its ability to finely control the stereochemical outcome of reactions while producing minimal amounts of waste and exhibiting an excellent resource-balance^2^. Early efforts in biocatalysis predominantly relied on natural enzymes, which often posed limitations in terms of scope and versatility. However, with the advent of directed evolution, a revolutionary opportunity arose to tailor native enzymes to targeted applications. This breakthrough facilitated the enhancement of enzyme activity for both native and novel substrates and the optimization of stereoselectivity and adaptability to diverse reaction environments^3,4^. In addition to these experimental advancements, computational tools offered valuable assistance in the engineering and analysis of experimental-directed evolution workflows The integration of computational methods into the biocatalysis toolkit represented a significant milestone, promising to expedite the development of next-generation biocatalysts with enhanced performance characteristics^5^. Moreover, the use of biocatalysts for synthetic purposes has contributed significantly to the transition of the chemical and pharmaceutical industries towards cleaner and metal-free production processes^6^. Most of these approaches relied either on the identification of a promiscuous activity in a naturally occurring protein or used natural protein scaffolds as starting points. In both situations the starting points are likely compromised by evolutionary effects^7^, a limitation that can be overcome by using de novo proteins^8–11^.

Custom designed de novo enzymes can become an alternative to natural starting points as recently highlighted by severl publications reporting on highly active luciferases, heme-dependent (oxidation) enzymes, xylanases, P450 enzymes and more^8,12–15^. A common feature of these studies is their high specificity for the targeted reaction and substrates. As a result, the current paradigm for designing efficient biocatalysts is to design entirely new proteins for each new substrate and reaction, often in small de novo scaffolds, which tend to be not very tolerant against variations in their amino acid sequence^16,17^. While this approach is valuable it could be sped up dramatically, if stable and highly engineerable scaffolds to catalyze envisioned reactions with or without cofactor binding are available. To achieve this, we need to address functional design and selection for activity within a single, highly stable protein fold, able to withstand destabilizing amino acid changes. This also offers the possibility to apply de novo biocatalysts for whole-cell biotransformations, an approach that streamlines applications for synthesis purposes as it bypasses costly and time-consuming purification steps.

Here, we present a highly versatile protein scaffold which acts as an efficient whole cell biocatalyst, designed by finely sampling helical arrangements of long straight alpha-helices using parametric design^18^. In addition to finding optimal packing solutions for an anti-parallel six helix barrel with high thermostability, we investigate placement of a potent catalytic array along the central channel of the design to probe its resistance to high mutational loads and exemplify its versatility as a re- usable asymmetric reaction chamber by designing nine variants. Biochemical and biophysical characterization as well as an experimentally determined structure allowed us to test several of the above-described features that are prerequisites for a universally applicable de novo protein backbone for biotransformations.

### Parametric design allows for ultrafine backbone geometry optimization

We set out to design a multifunctional de novo protein chassis, which can be used as an asymmetric reaction chamber. To do so, we aimed at designing anti-parallel six-helical barrels accommodating ligand or substrate accessible channels, starting from the design principles elucidated during the design of four-helix bundles^18^. To identify stable arrangements, we extensively sampled super-helical coiling, radius and the translation along the central axis of the individual helices. Loops to connect the individual helices were kept short and idealized. This resulted in lowest energy sequences for totally straight helical arrangements with 18-residues repeating units and an average diameter of 15.7 Å (Cα-Cα of opposing helices, Figure 1A). In contrast to smaller helical bundles, the large helical barrels do not allow for tightly packed cores. Thus, the biggest contribution to structural integrity is coming from neighboring anti-parallel helices. Naturally occurring helical barrels with similar numbers of anti-parallel helices typically feature oligomerization of smaller, stable entities or stabilizing domains on each side of the helix barrel due to the lack of a stably packed core^19^, which makes this approach very challenging from a design perspective. The backbones of these designs were generated as described previously^18^ and by using the parametric design code as implemented into Rosetta^20^. The straight helices and overall configurations made it possible to install salt-bridges across the barrel at two distinct positions, thereby, enhancing the folding and stability of the generated designs (Figure 1A).

**Figure 1.**
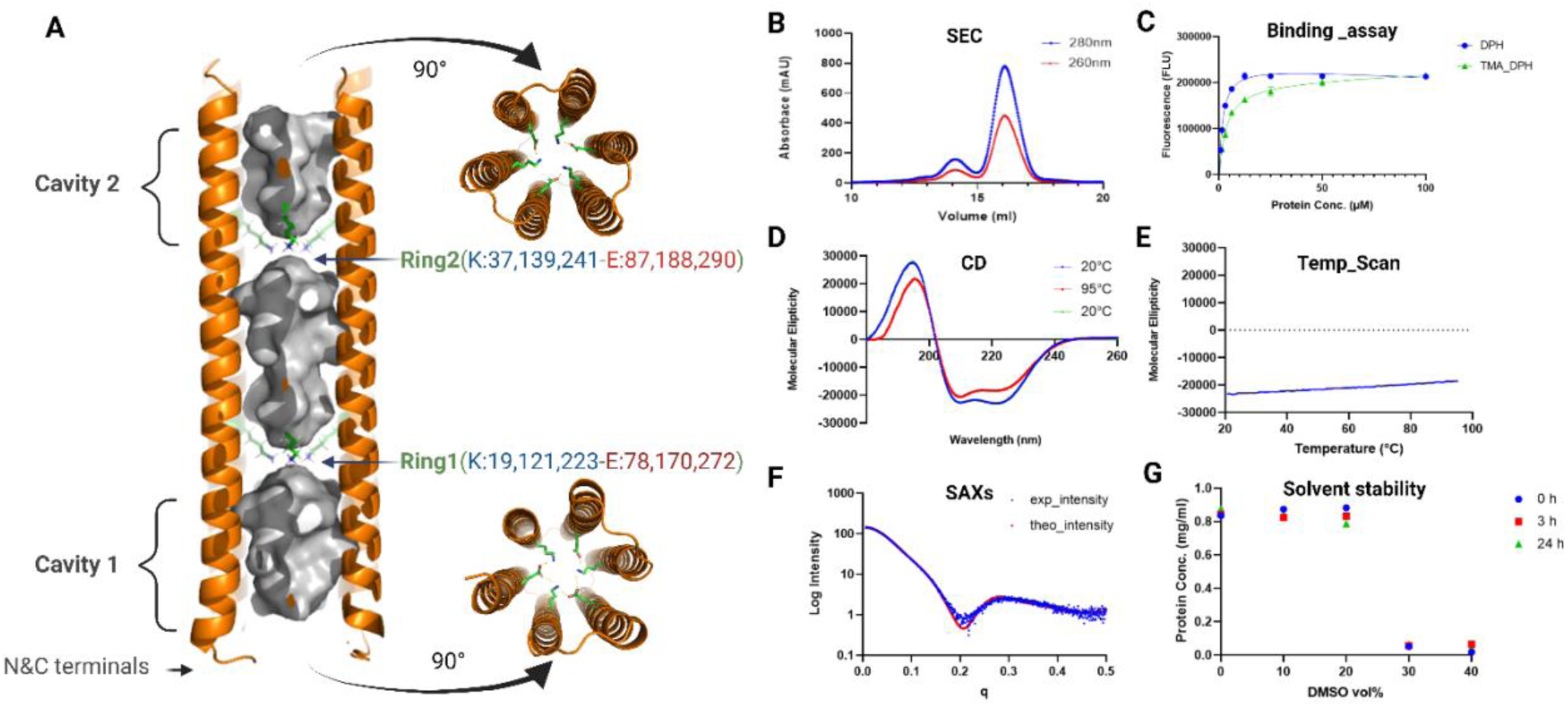
Characterization of 6H5L design **A)** Structure representation consists of a six-helix barrel, with two rings introduced into the hydrophobic channel, creating two cavities, each consists of three lysine (K) and three glutamate (E) residues connected by electrostatic interactions, contributing to structural stability and enhanced catalytic efficiency. **B)** Size exclusion chromatography (SEC) of 6H5L desired monomer fraction as the highest peak at the expected retention volume with good separation from the dimer form. **C)** Saturation binding curves of different 6H5L protein concentrations with 1μM constant concentration of 1,6-diphenylhexatriene (DPH) or 1-(4-trimethylammoniumphenyl)-6-phenyl-1,3,5- hexatriene *p*-toluenesulfonate (TMA-DPH). This result indicates that the protein folds in a barrel shape giving rise to a hydrophobic channel. **D)** Circular dichroism (CD) spectrum confirmed an all alpha-helical fold. **E)** CD Temperature scan exhibited high thermal stability with only small changes until 95°C with ideal refolding by cooling back to 20°C. **F)** SAXS solution scattering plot showing good fitting between the theoretical (red) and measured (blue dots) curves with chi-square of 2.2. Curve fitting was executed using the FoXS online server. **G**) Solvent stability was measured by determining the soluble protein concentration using absorbance at 280 nm, plotted against DMSO concentration at three time points (0h in red, 3h in light blue, and 24h in gray), indicating protein stability over time up to 20 vol% DMSO.

We generated eleven versions of such designs with different twists and repeating units, however, only one of the designs – 6H5L – could be produced in high amounts in the wet lab and exhibited an all-helical circular dichroism (CD) signal and high thermostability (Figure 1C&D).

Interestingly, during combinatorial sequence design calculations, we found that the frequency of alanine incorporation at helical interface positions strongly depended on the set supercoil and individual helical coiling parameters, with some designs being overpopulated with this residue type, even though no obvious geometry or space restrictions were apparent. Therefore, we adjusted the Rosetta reference energies^21^ to achieve the desired residue identity balance. All the herein described designs were computed prior to the advent of machine learning based structure prediction methods and thus could not be tested in this respect prior to ordering synthetic genes. However, we and others recently found that AlphaFold2^22,23^ could accurately predict the structures of designed helical barrels from single-sequence input^11^. Therefore, we checked 6H5L and all its variant sequences subsequently on their prediction quality (suppl Figure S4).

### Characterization of the parametrically designed helix bundles in *E. coli*

Production of the parametrically designed protein 6H5L and all its variants yielded soluble protein after production in *E. coli.* The proteins were purified using affinity and size-exclusion chromatography, yielding nearly identical SEC chromatogram profiles for 6H5L and its variants, featuring a prominent absorbance peak at 280 nm with the expected retention volume for a protein of this size. In addition, the chromatograms displayed a second minor peak, which corresponded to a dimeric form of our designs. However, separation between the monomeric and dimeric form could be achieved (Figure 1B). The pooled and concentrated fractions from the monomer peak were confirmed to contain highly pure protein (Suppl Figure S2A&B).

To confirm folding and the secondary structure content of our designs, we recorded CD spectra for both pure 6H5L (Figure 1D) and its variants (Suppl Table S6). All recorded spectra exhibited a characteristic α-helical profile. Notably, these structures exhibit very high thermal stability as exemplified by only a minimal loss of signal (222nm) during temperature ramps up to 95 °C (Figure 1C, 1D & Suppl Table S6) and refolding after cooling back down to 20°C. This highlights the robustness of the designs, even when introducing mutations in the core for active site redesign (all single lysine and active-site redesigned variants) or on the surface for improved crystal formation (FKQ variant). A summary of all CD spectra from all variants can be found in Supplementary Table S6. To assess barrel formation, we used previously described binding experiments of fluorescent molecules.^24^ One of the benefits of this assay is that it can be performed in a semi-high throughput manner using 96-well plates. All designs saturation binding behavior, which could be fitted to a one-site total binding model for calculating dissociation constants (*K_D_*). The observed affinities are in the micromolar range with specific *K*_D_s between 1.1 to 4.5 µM (Figure 2B and Supplementary Table S6). These results corroborate the designed barrel-like structures for all designs. To further investigate the correct folding and dynamics of our designed proteins, we employed SAXS measurements. The obtained results show good fits between the measured and calculated scattering profiles, characterized by low chi² values, particularly within the Guinier region of the logarithmic plot (Figure 1F and Suppl Table S6). All obtained fits indicate that the designs exhibit the expected behavior in solution, consistent with the AlphaFold2 predicted models^23,25^.

**Figure 2.**
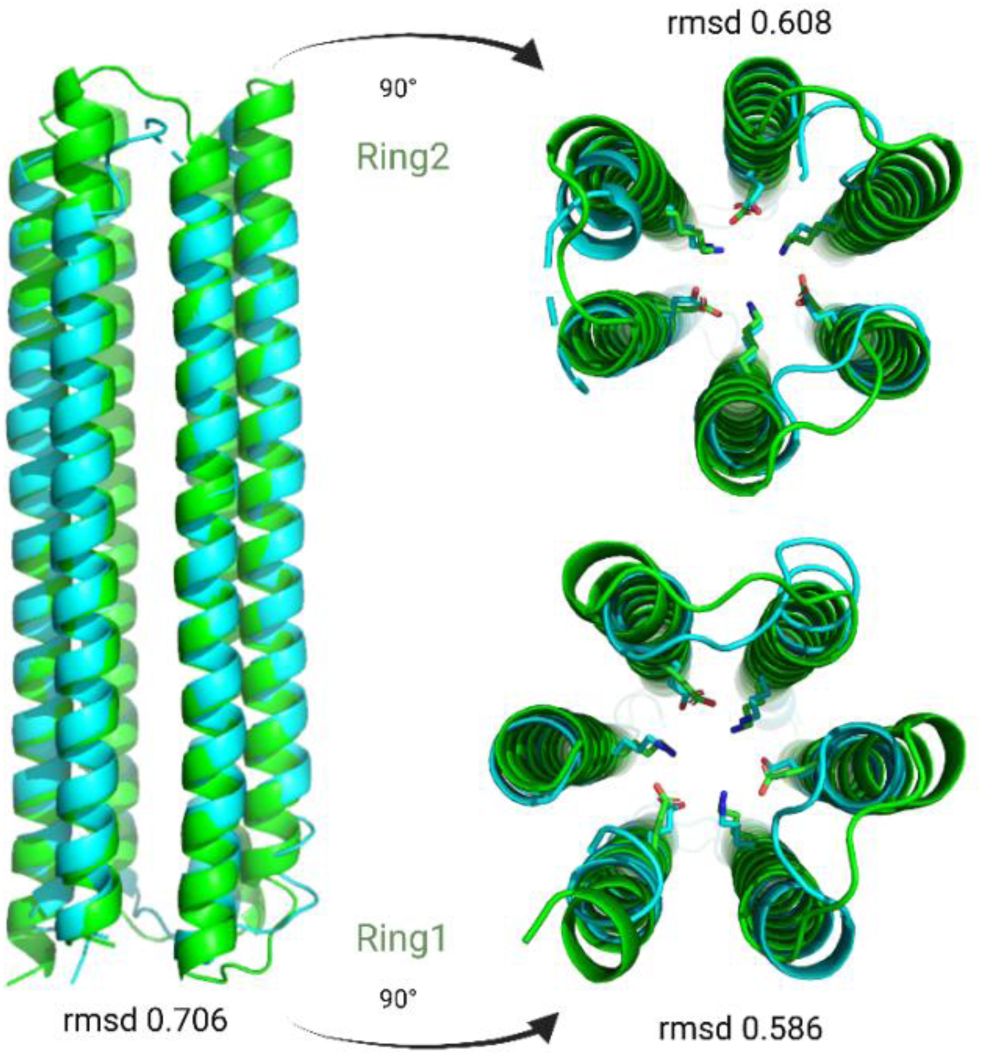
Superposition of parametrically designed 6H5L (in green) and the experimental crystal structure of 6H5L_FKQ (in cyan), highlighting the high overall similarity between model and structure. Residues constituting the two stabilizing electrostatic rings in the center of the structure adopt highly similar geometries in the design and the experimental structure.

An important feature of a practically applicable biocatalyst is its stability and activity in organic (co)solvent. We assessed the stability of 6H5L in up to 30 vol% DMSO over time points of 0, 3, and 24 hours. Notable instability was observed only in concentrations of 30% DMSO and higher, highlighting the proteins high solvent tolerance (Figure 1G).

### Protein crystallization and structure determination

Initial crystallization of the original 6H5L sequence resulted in the formation of 2D crystals only and ultimately did not produce high enough diffraction quality to determine an experimental structure. However, protein modification to enhance crystal contact formation by exchanging a total of 15 lysine residues on the equivalent helical wheel *f* position with glutamine (6H5L_FKQ variant), lowered the pI from 8.99 to 5.7 and resulted in well-ordered three-dimensional crystals diffracting up to 2.8 Å. To ensure that this extensive residue exchange did not interfere with correct folding, we used AlphaFold2 to predict the 6H5L_FKQ structure and found that it closely matched the original 6H5L, with an RMSD of 0.17 across all atoms.

The 2.8 Å structure of 6H5L_FKQ shows an overall high degree of similarity to that of the design model (Figure 2) with the predicted straight helices, no supercoil twist and the 18-residue repeating unit. The packing differences within individual repeats between design and structure are minimal with an Cα-atom RMSD of ∼0.7 Å over the core repeating units. This structure is another example of a computationally designed helix bundle that deviates from perfect supercoil geometry, further corroborating that low energy structures for monomeric antiparallel helix bundles are not strictly confined to the space spanned by the Crick parameterization, particularly not near the turns.

### Exploring the catalytic potential of the *de novo* designed structure

The design model of 6H5L, confirmed by crystallography, presents an open, straight six-helix barrel with a hydrophobic channel, highly effective at binding hydrophobic aromatic molecules, as proven by DPH and TMA-DPH fluorescence assays Figure 1C). Crystallographic analysis also identified two hydrophobic cavities within the barrel, each accommodating three lysine residues, electrostatically coupled with three glutamates, forming two identical electrostatic rings. We reasoned that we could exploit the nucleophilicity of the central lysine residues to catalyze enzymatic reactions. Experimental characterization of variants, with only a single lysine in the central channel and the remaining five replaced by arginines, showed similar behavior to the original 6H5L design. Having a potentially nucleophilic lysine present, we opted to assay the designs for the retro-aldol reaction, utilizing methodol **3** as a substrate (Figure 3A).

**Figure 3.**
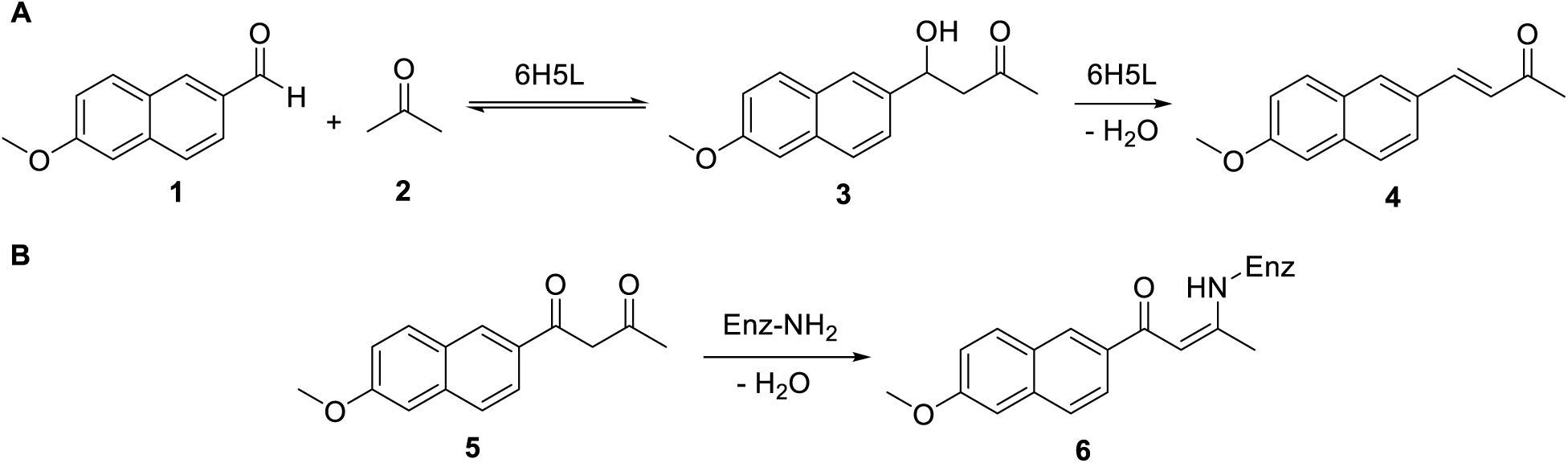
Model reactions. **A)** Reversable aldol reaction of 6-methoxy-2-naphthaldehyde **1** and acetone **2** producing 4-hydroxy-4-(6-methoxy-2-naphthyl)-2-butanone (methodol, **3**) and the α,b-unsaturated ketone **4** upon dehydration. **B)** 1,3-diketone **5** as inhibitor forming the covalent enamine adduct **6** with the catalytically active lysine.

As a proxy to assess the ability of the original 6H5L design and its variants to catalyze retro-aldol reaction through an active nucleophilic lysine, we conducted an inhibition reaction (Figure 3B) using the corresponding 1,3-diketone derivative **5** (1-(6-methoxy-2-naphthalenyl)-1,3- butanedione) of methodol **2**. This derivative is expected to covalently react exclusively with the catalytically active lysine residue (H2N-Enz) through a Schiff-base intermediate, forming a stable vinylogous amide **6**.

After the reaction of **5** (Figure 3B) with 6H5L and its six single lysine variants, the resulting UV-Vis absorbance spectra show a new peak at 340 nm compared to the free protein (Figure 4A). This indicates the successful formation of a covalent adduct, underlining the capability of these proteins to interact with a carbonyl group and thus potentially to catalyze a retro-aldol reaction. The intensity of the adduct absorbance peak increases with an increasing diketone-to-protein ratio (1:1, 5, and 10), indicating more diketone binding with higher concentration (Figure 4A). The intensity of the peak of the adduct between **5** and 6H5L is approximately double that of the peaks of the addict from its single lysine variants (Figure 5A). Given the distribution of the 6 lysine in two cavities, this suggests one single lysine is reactive per binding pocket.

**Figure 4.**
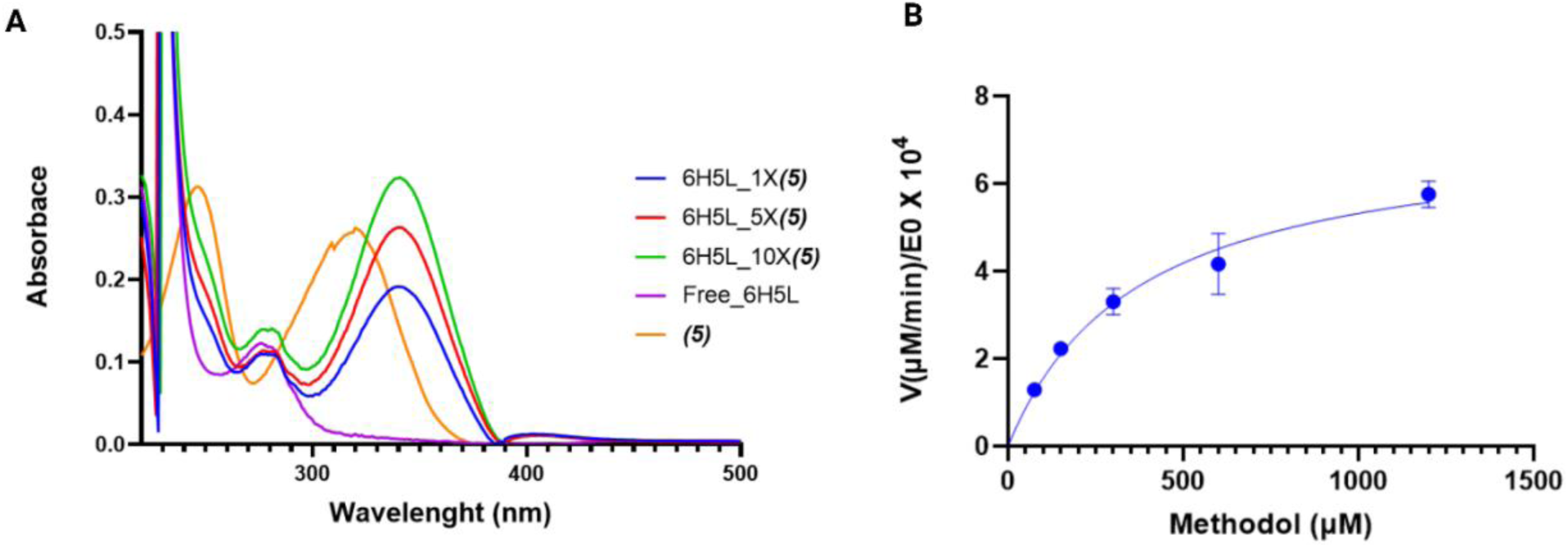
**A)** UV-VIS absorbance spectrum of a free 6H5L (27 µM) and labeled with varying concentrations of **5** (1X, 5X, and 10X) showing an additional peak at 340 nm compared to the free protein. Higher concentrations of **5** led to increased peak intensity. **B)** Retro-aldolase Michaelis–Menten plots with methodol as a substrate of 6H5L design. E0, total enzyme concentration; V, initial reaction velocity.

**Figure 5.**
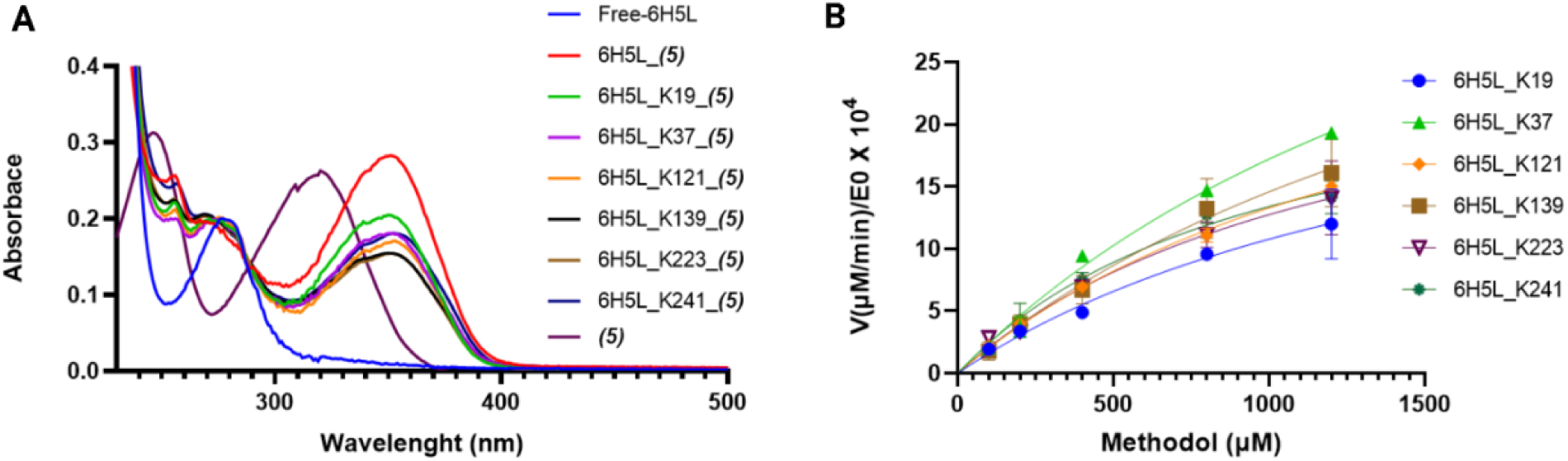
**A)** The UV-VIS absorbance spectrum of diketone-labeled 6H5L and all its single lysine variants (all at 27 µM) reacted with 270 µM diketone **5** revealed an additional vinylogous amide peak at 340 nm compared to the free protein. The vinylogous amide peak of 6H5L exhibited nearly double the intensity compared to its variants. **B)** Retro-aldolase Michaelis–Menten plot of all 6H5L single lysine variants with *rac-*methodol as a substrate. E0, total enzyme concentration; V, initial reaction velocity. Notably, all variants exhibited significant enzymatic activity.

To exclude spurious activity from surface lysine residues and to confirm specific activity through the lysine residues in the central channel, we performed competition assays between the diketone inhibitor **5** and DPH fluorescence dye. The first competition assay showed a decrease in fluorescence intensity for the diketone-labeled 6H5L in comparison to the free 6H5L protein (Suppl Figure S5A). This decrease persisted even at higher diketone inhibitor concentrations and provides evidence of diketone binding within the hydrophobic channel of 6H5L, thereby effectively blocking the central channel and preventing the binding of DPH. The reverse way of the competition assay, involving the titration of diketone inhibitor into 6H5L previously saturated with DPH, demonstrated a progressive decrease in fluorescence intensity with the addition of more diketone inhibitor resulting in an inhibition constant (Ki) of 0.27 mM (Suppl Figure S5B). These findings corroborate the binding of the diketone inhibitor to the lysine residues within the central channel.

To test for retro-aldolase activity, a well-established fluorescence-based method to measure the formation of a naphthaldehyde derivative (6-methoxy-2-naphthaldehyde) **1** as product from racemic methodol **3** as a substrate, was used (Figure 3A)^26^. The original 6H5L design demonstrated noticeable retro-aldolase activity with a second-order rate constant, *k*_cat_*/K*_M_, of 1.96 M^−1^ min^−1^ (Figure 5B. and Table 1). This rate is comparable to some of the first *de novo* retro- aldolases^27^. This underscores its potential as a promising retro aldolase candidate by further improvement. An important validation experiment for this is checking the pH dependence of the monitored reaction. We found that the activity of 6H5L decreased at lower pH levels in the reaction media but remained significantly higher compared to the knockout version (6H5L_6-R), which lacks any lysine residues inside the two binding pockets. The knockout version (6H5L_6-R) exhibited almost no activity at pH 6, whereas 6H5L remained visibly active under the same conditions (Suppl Figure S6). This illustrates the nucleophilic activity of the lysine residues in the hydrophobic channel while protonated lysine residues on the surface at lower pH remain inactive.

**Table 1.** Kinetic parameters of retro-aldolase for active site redesigned variants of 6H5L and its initial input designs, in comparison to a variety of initially computationally designed enzymes (assessed using the racemic methodol).

| Protein | $K_m$<br>($\mu\text{M}$) | $k_{cat}$<br>($\times 10^{-4} \text{ min}^{-1}$) | $k_{cat}/K_M$<br>($\text{M}^{-1} \text{ min}^{-1}$) |
| --- | --- | --- | --- |
| 6H5L | 371.4±62 | 7.3±0.1 | 1.96 |
| 6H5L_K223 | 967 ±211 | 9.5 ±1.1 | 0.98 |
| 6H5L_RA2 | 1550 ±225 | 45.0 ±4.2 | 2.90 |
| 6H5L_RA1 | 1037 ±128 | 74.00 ±5.3 | 7.14 |
| 6H5L_Top | 453.2 ±49.31 | 95.0±4.5 | 20.95 |
| RA95<sup>32</sup> | 530 | 60 | 11 |
| RAβb-9<sup>33</sup> | -- | -- | 17 |

### Single lysine and knockout variants

To identify the most catalytically proficient lysine among the six within the hydrophobic channel, we designed six single-lysine variants (6H5L_K19, 6H5L_K37, 6H5L_K121, 6H5L_K139, 6H5L_K223, 6H5L_K241). Each variant features one lysine in the hydrophobic channel, with the other five lysine residues being replaced by arginine to maintain the electrostatic interactions that stabilize the barrel structure (Suppl Figure S3). Additionally, we used the knockout variant 6H5L_6-R, which lacks any lysine residues within the two binding pockets as a negative control. All six single lysine variants exhibited measurable activity over time, albeit with lower activity than the original 6H5L (Figure. 5B and Table 1). This suggests the possibility of both cavities being active and highlights the influence proximity effects of nearby lysine residues in lowering the *pKa* of the nucleophilic lysine.

### Placing of a catalytic tetrad enhances the activity of the designed retro-aldolase

We aimed to enhance retro-aldolase activity by introducing the catalytic array of the well-studied laboratory evolved and highly efficient retro-aldolase RA95.5-8F ^28^ into 6H5L. This involved incorporation of two tyrosine residues, one glutamine, and a catalytic lysine (KYQY tetrad) to form a fully connected hydrogen bond network. Initially, we selected the single lysine variant 6H5L_K223 as our starting point and used Foldit^29^ as an interface to Rosetta to redesign three residues near this lysine to the corresponding tyrosines and glutamine. The resulting model was repacked and minimized, which resulted in the design of the 6H5L_RA2 (Figure 6A). In a second approach, we used RosettaMatch^30^ for a comprehensive search for geometric matches of 1,3- diketone **5** (1-(6-methoxy-2-naphthalenyl)-1,3-butanedione) and the residues form the evolved active site of RA95.5-8F within our parent 6H5L design model. Initially, we confined this search to the central lysine resides and their immediate surroundings, which led to the design 6H5L_RA1. Furthermore, matching across the entire scaffold, led to the design variant 6H5L_Top (Figure 6A). Combinatorial sequence design in Rosetta^31^ was used to optimize the amino acids surrounding the cavities hosting the ligand. The best scoring design was superimposed onto the best-ranked AF2 predicted structure of the same sequence^2,3^. The two designs 6H5L_RA1 and 6H5L_Top (Figure 6A) exhibited the lowest rmsd were selected and ordered as synthetic genes for structural and functional characterization. All active site redesign versions were successfully produced and characterized using our standard methods (Supplementary Table S6). The rational (6H5L_RA2) and computational (6H5L_RA1) redesigns of the active site exhibited a 3-fold and 3.5-fold increase in catalytic efficiency compared to the single lysine variant 6H5L_K223 and the original 6H5L design, respectively (Suppl Figure. S7A and S7B, Table 1). To confirm that the increase in catalytic efficiency arises from the newly installed active site and redesigned cavity, knockout variants 6H5L_RA2_K223A and 6H5L_RA1_K37A were tested. The results showed a dramatic decrease in activity, reverting almost to the levels observed in the initial input designs (Suppl Figure S7C&D and Table S4). Notably, the 6H5L_Top version exhibited a remarkable 10-fold activity improvement achieved by repositioning the active site higher above the hydrophobic channel (Figure. 6A, 6B and Table 1). This strategic adjustment significantly enhanced the catalytic efficiency nearly twofold in the second-order rate constant (*k_cat_/K_M_*, of 20.95 M^−1^ min^−1)^ compared to the TIM-barrel computationally designed retro-aldolase RA95 (*k_cat_/K_M_*, of 11 M^−1^ min^−1^)^32^ and slightly higher than RAβb-9 *de novo* β-barrel scaffold (*k_cat_/K_M_*, of 17 M^−1^ min^−1^)^33^, addressing one of the most challenging aspects in enzyme evolution^34^.

**Figure 6.**
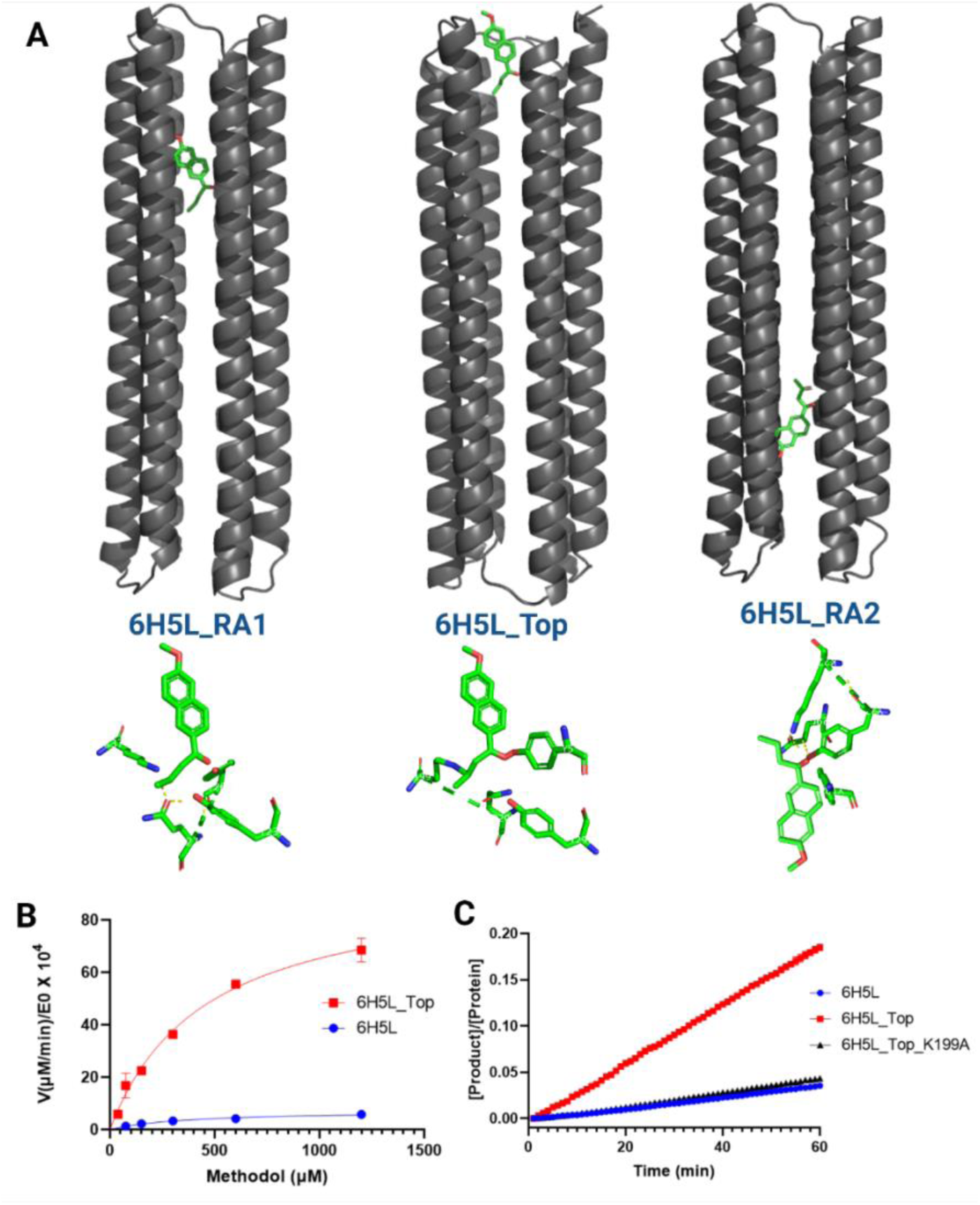
**A)** 6H5L active site redesign variants, including methodol **3** substrate, are positioned in the redesigned pocket at different locations within the hydrophobic channel. **B)** Retro-aldolase Michaelis–Menten plots with methodol **3** as a substrate of 6H5L_Top (in red) active site redesigned variant compared to the initial 6H5L (in blue) activity of the input designs. E0, total enzyme concentration; V, initial reaction velocity. 6H5L_Top variant: 10-fold increase compared to 6H5L (Table 3). **C)** Retro-aldolase progress curves of 6H5L_Top (in red) active site redesigned variant of 6H5L compared to and its initial 6H5L (in blue) input design and 6H5L_Top_K199A (in green) knockout version.

To confirm that the increase in catalytic efficiency originates from the expected lysine at position 199, a knockout version was generated and tested. 6H5L_Top_K199A showed a roughly 10-fold activity decrease, which is comparable to the original 6H5L design (Figure 6C and Suppl Table S4).

The biophysical characterization and activity results, it’s evident that the targeted changes to the initial 6H5L backbone/sequence, especially within the hydrophobic channel, have led to improved activity without significantly influencing the overall structure (Supplementary Table S6).

### Two-step reaction (Aldol/Retro-aldol and dehydration)

The aldol and retro-aldol (Figure 3A) catalytic activities of 6H5L and 6H5L_Top were evaluated by HPLC and GC/MS to quantify conversion and enantioselectivity over a 24 h period. In the aldol reaction (Figure 1A) between 6-methoxy-2-naphthaldehyde **1** (5 mM) and acetone **2** (10% v/v), both proteins exhibited clear catalytic activity, consuming 28% and 41% of **1**, respectively (Figure 7A). Beyond C–C bond formation yielding Methodol **3**, both variants promoted dehydration to the α,β-unsaturated ketone **4**, which emerged as the predominant product (1.8 and 2.2 mM). Only minor amounts of racemic **3** were detected, with enantiomeric excess values of 10.4% and 7.5% (Figure 7B, Suppl Figure S9A). In the retro-aldol reaction with racemic **3** (1 mM), 6H5L displayed non-enantioselective with substrate consumption (10.3%), whereas 6H5L_Top preferentially consumed the *(S)-* and *(R)-*enantiomers (15.5% vs. 13.7%; ee = –7.9%). Both proteins produced aldehyde **1** at 18 and 106 µM, respectively, and generated the dehydrated product **4** (73 and 160 µM; Suppl Figure S8 & S9B). Taken together, the data indicates that both designs are active, with 6H5L_Top giving higher conversion and favoring aldehyde formation over dehydration compared to 6H5L in the retro-aldol reaction.

**Figure 7.**
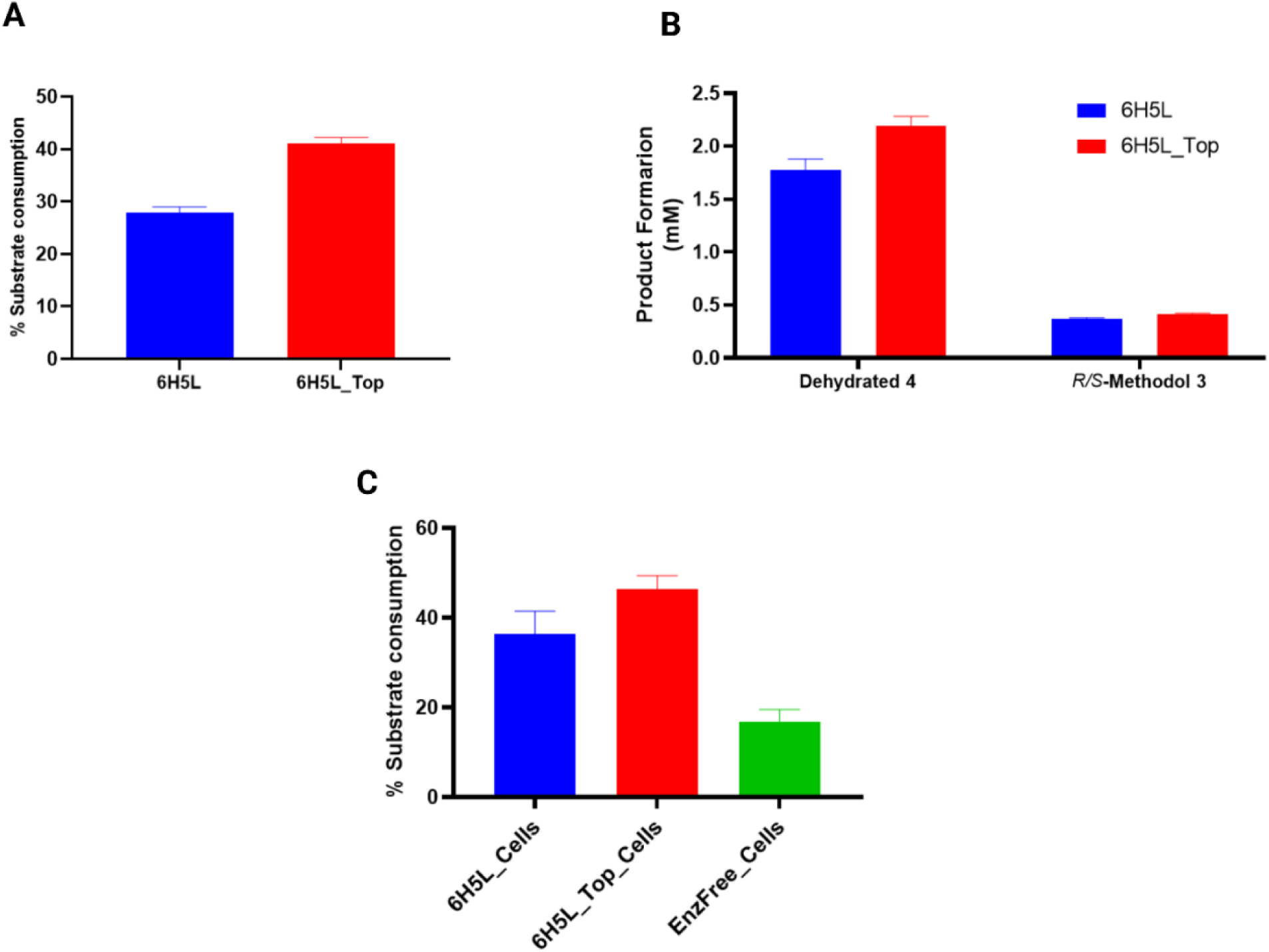
Aldol reaction catalyzed by 100 µM 6H5L (blue) and 6H5L_Top (red) using 5 mM 6-methoxy-2-naphthaldehyde **1** and 10% acetone **2** in 50 mM sodium phosphate buffer (pH 8) containing 150 mM NaCl and 20% acetonitrile at 30 °C**. A), B)** Substrate consumption (%) of **1** and Products formation of (*R&S*)-Methodol **3** and its dehydrated derivative **4** using 100 µM purified enzyme. **C)** Substrate consumption (%) in whole-cell catalysis compared with enzyme-free cells (EnzFree_Cells) using 5 mM **1** and 5% **2** in the same buffer containing 10% acetonitrile (pH 7.5).

Reducing acetonitrile (from 20% to 1%) and acetone **2** (from 10% to 5%) in the aldol reaction (Figure 1A) decreased activity relative to the previous reaction conditions, with substrate **1** (1 mM) consumption of 9.2% for 6H5L and 6.3% for 6H5L_Top. The ratio of dehydrated product **4** formation declined compared to aldehyde **3**, which also exhibited opposite enantiomeric preferences (ee *R/S* = +28.5% for 6H5L, –7.6% for 6H5L_Top; Suppl Figure S10), indicating that co-solvent and substrate levels can be adjusted to fine-tune conversion, enantioselectivity, and product distribution.

We assessed the long-term stability of 6H5L and 6H5L_Top after storage at 4 °C for over three months; both enzymes retained aldol activity between 1 (5 mM) and 2 (5% v/v), with substrate conversions of 7.3% and 6.2%, respectively, comparable to freshly prepared samples under identical conditions (Suppl Figure S11).

### A practically useful de novo enzyme - whole cell biotransformations

To be applicable to industrial applications, most biocatalysts are used in a whole cell fashion, which means that they can be produced in large fermenters and used directly in reaction mixtures, without the need for further purification. To showcase the high versatility of our designs, we tested 6H5L and the more active variant 6H5L_Top as whole cell biocatalysts for the aldol reaction (Figure 3A) of 6-methoxy-2-naphthaldehyde **1** (5 mM) and acetone **2** (5% v/v). Both designs exhibited significant activity, with substrate consumption around 36% and 46%, respectively, while enzyme-free cells showed only 16% conversion (Figure 7C). The aldol product was found to undergo dehydration to the corresponding α,b-unsaturated ketone **4**, and control reactions indicated that this dehydration was in the most part catalyzed by the protein (*vide infra*).

Additionally, 6H5L overexpressing cells catalyzing the retro-aldol reaction (Figure 3A) on 5 mM of **3**, showed 35% consumption of *rac*-methodol in a non-enantioselective behavior. A similar trend was observed with cells expressing the construct 6H5L_Top (36% substrate consumption, Suppl Figure S12). Notably, the catalyst preparation was compatible with the organic solvent acetonitrile, up to 20 vol%.

In line with the observed formation of the aldol condensation product **4** through dehydration as major product during the reaction between **1** and **2**, we could detect the formation of **4** during the retro-aldol reaction with **3** and confirmed that this reaction was catalyzed by the over expressing cells. The significant aldolase and retro-aldolase activity in whole cells, both from the original 6H5L design and the 6H5L_Top active site redesign variant, highlights the potential of this de novo helix barrel to be used as a highly engineerable reaction chamber. Combined with its high thermodynamic stability and solvent tolerance to 20 vol% this design established itself as a promising candidate for optimization and utilization in green biotechnology applications.

### Conclusions

Enzyme design efforts currently rely on designing and screening up to hundreds of hypothetical enzymes to produce a viable catalyst. While this is a highly efficient strategy, it lacks the ease that many of the orthogonal approaches, like organocatalysis, exhibit - to quickly and rationally engineer and repurpose catalysts for envisioned reactions. One way of gaining quicker access to products of interest and using the exquisite asymmetric environment that a protein provides, is via a highly versatile and stable protein chassis with a high mutational tolerance. Our study constitutes a significant step forward in this direction. Using the highly accurate backbone generation via parametric design, we demonstrated that such a scaffold for holding catalytic centers in an asymmetric environment can be made and yields moderately efficient enzymes while screening as little as nine variants. The ability to position catalytic sites at different locations within the same scaffold further provides flexibility for engineering catalytic function. While the most active design does not show an improvement over previous in-silico designs, its high thermostability and solvent tolerance allows it to be used as an efficient whole cell biocatalyst. This highlights the compromise between ease of engineering and high-rate accelerations, usually only achieved through lengthy laboratory evolution campaigns. However, the ability to scaffold catalytic centers in a robust protein chassis with high accuracy now allows the type of experiments that can elucidate such structure- and function relationships quickly, helping to build models for synthetic strategies in biomolecules.

Thorough biochemical characterization allowed us to identify practically useful enzymes that exhibited high stability and moderate turnover in whole cell biotransformations. For generalized models of proteins being used as asymmetric reaction chambers, datasets like this will be needed for a diverse set of chemical transformations. In addition, the ability to design protein scaffolds like ours could foster the creation of a set of asymmetric environments that can be highly engineered to be used as versatile biocatalysts. Much like the streptavidin system has been used to anchor artificial cofactors to an asymmetric environment and thereby fostered the development of new synthesis routes, our approach could drive the development of methods for reusability of de novo designed functional proteins.

Finally, whole cell biotransformations have played a pivotal role in biocatalysis campaigns to create synthetic routes to new pharmaceuticals and complex fine chemicals. Here, we established a protein scaffold which shows great promise in being generally applicable for biotransformations. More broadly, the defined architecture and tunable active-site positioning of these scaffolds could be extended to other applications where stable and engineerable protein environments are required, such as biosensing and biosynthetic systems.

## Methods

### Parametric design of six-helix barrels

Parametric design was described in detail before.^18^ Briefly, envisioned helical barrel geometries were generated using the Crick coiled coil generating equations, as described previously^18^ or using the bundle grid sampler, as implemented in Rosetta^20^ for specified parameter ranges. The helix barrel was built from an asymmetric input of an up- and a down-helix arranged around a threefold (C3) symmetry axis. Rosetta was used to regularize the backbone geometry by minimization in torsion and cartesian space and low energy sequences were designed onto the backbones by Monte Carlo Rotamer optimization, followed by sidechain minimization, first with the repulsive interactions damped and then undamped (a detailed input xml file for RosettaScripts to carry out minimization and design is provided in the Supplemental material). Loops were built onto low energy models using RosettaRemodel^20^, and the geometry again regularized by cartesian space minimization. Sampling of backbone parameters and sequence design was performed until low energy fully connected designs with near-ideal backbone geometry and few alanine and glycine residues were found.

### Active site redesign

In a semi-rational approach, we started from the single-lysine variant 6H5L_K223 (Suppl Figure. S3) and used Foldit^29^ as an interface to Rosetta^20^ to redesigned three residues close in space of K223 by mutating two of them to tyrosine, and one to asparagine to install an active site similar to the catalytic tetrad observed in RA95.5-8F. The resulting model was repacked, minimized, and relaxed with constraints to ensure the best possible alignment with the catalytic array from RA95.5- 8F. This resulted in the design 6H5L_RA2 (Figure 6A).

In a subsequent approach, we used RosettaMatch^30^ to search for geometric matches between 1,3-diketone **5** and the catalytic array as found in RA95.5-8F^28^. Geometric constraints were extracted from the inhibitor bound crystal structure of RA95.5-8F to create a constraints file for matching within 6H5L (match_cstfile is provided in the Supplemental Material). A focused search around the six lysine residues in the two electrostatic rings inside the barrel lumen, led to the design 6H5L_RA1 (Figure 6A). Expanding matching to include all positions in the scaffold resulted in optimal substrate placement and catalytic geometry close to the entrance of the barrel and the variant called 6H5L_Top (Figure 6A). After matching, Rosetta^20^ combinatorial sequence design was used to optimize amino acids around the substrate binding site (a detailed input XML file for RosettaScripts is provided in the Supplemental Material), with the best-scoring designs matching their AlphaFold2^23,25^ predicted structures very closely.

### Protein production and purification

All individual protein variants were ordered as gene fragments and cloned using Gibson assembly^35^ into the pEThTEV plasmid (Suppl Figure. S1), which includes an N-terminal His-Tag with a TEV cleavage site. The successfully confirmed cloned genes were transformed into *E. coli* BL21 Star (DE3) competent cells for protein production. Single colonies were used to inoculate 20 mL LB pre-culture medium containing 50 mg/L Kanamycin and incubated overnight at 37 °C and 180 rpm. One liter of TB main culture containing 100 mg/L kanamycin was inoculated by adding 10 mL of pre-culture and incubated at 37 °C,180 rpm until an OD600 around 0.6 was reached. For protein induction, 1 mM isopropyl-β-D-thiogalactopyranoside (IPTG) was added to the main culture and incubated overnight at 18 °C and 180 rpm. The cells were harvested by centrifugation at 4 °C / 4500 rpm for 30 min and stored at -20 °C before lysis. The cells were lysed by sonication for 10 min under cooling in lysis buffer (20 mM sodium phosphate pH 7.4, 500 mM NaCl, 10 mM imidazole, and 10 mM PMSF). The lysate product was centrifuged at 18,000 rpm for 45 min to isolate the cell debris and the insoluble protein.

The protein variants were purified by Ni-NTA affinity chromatography and size-exclusion chromatography (SEC), including His-Tag cleavage using TEV protease for further biochemical characterization as follows. The cleared lysate was loaded onto the lysis buffer-equilibrated Ni-NTA column and washed with a wash buffer containing the same buffer but with 30 mM imidazole. The target protein was eluted from the column using elution buffer (20 mM sodium phosphate pH 7.4, 500 mM NaCl, 250 mM imidazole), concentrated and buffer exchanged to 20 mM Tris-HCl, 150 mM NaCl, pH 8. Tag cleavage was performed by adding 0.1 mg/mL TEV protease to the solution at a protein concentration of 1 mg/mL and incubating overnight at room temperature. The cleaved protein was separated from the mixture by loading it again onto a Ni-NTA column and collecting the flowthrough. Size-exclusion chromatography was performed as the second purification step, using Xtal buffer as the mobile phase over Superdex 200 column. SDS-PAGE gels were run to monitor the protein throughout all purification steps. The concentrations of the pure proteins were calculated by measuring the absorbance of the samples at a wavelength of 280 nm using the A280 measurement option of the NanoDrop™ 2000 for all further biochemical characterizations.

### Circular dichroism spectroscopy (CD)

For CD measurements, the pure proteins were run over Size-Exclusion Chromatography (SEC) once again using CD buffer (20 mM sodium phosphate and 150 mM sodium fluoride at pH 8.0) as the mobile phase. The eluted fractions were collected and concentrated to 1 mg/ml. Measurements were carried out using quartz capillaries (Helix Biomedical Accessories, Inc.) with a path length of 0.5 mm on a CD spectrometer (J-1500, JASCO, USA). Wavelength scan spectra were measured from 260 to 190 nm, and temperature scans were performed with incremental steps of 2°C from 20°C to 95°C and back to 20°C. The CD signals were corrected using a blank sample containing the same CD buffer composition.

### Protein solvent stability

The purified 6H5L protein was dissolved in a buffer of 20 mM Tris-HCl and 150 mM NaCl, pH 8, to a final concentration of 1 mg/ml, with DMSO concentrations varying from 0 vol% to 50 vol%. Before each measurement, the mixture was centrifuged for 5 minutes at 14,500 rpm and 4°C for clarification. UV-Vis absorbance spectra and protein concentrations were measured at different time intervals (0, 3, and 24 hours) to monitor the protein’s solubility under each condition.

### Ligand-binding

Ligand-binding assays were performed using 1,6-diphenylhexatriene (DPH) or 1-(4- trimethylammoniumphenyl)-6-phenyl-1,3,5-hexatriene p-toluenesulfonate (TMA-DPH) fluorescence days and all our designed helix bundle variance, The experiments maintained a constant ligand concentration of 1 µM, while the designed proteins concentrations varied from 0.78 or 1.6 up to 50 or 100 µM in 20 mM Tris, 150 mM NaCl, 1% DMSO, pH 8 buffer at 25 °C. After a 1-hour incubation with shaking at 300 rpm, fluorescence spectra were obtained from the top of the samples in a 96-well plate, using specific excitation and emission wavelengths (Table 5). The dissociation constant (K_D_) values were determined by fitting the data to a one-site_total binding model using GraphPad Prism 8. All the measurements were carried out in 96-Well Black flat bottom polystyrene non-binding surface microplates (Corning, Lowell, MA) using a BMG Labtech CLARIOstar PLUS Microplate Reader (Aylesbury, UK).

### Small-angle X-ray scattering (SAXS)

For SAXS measurements, each protein variant was run over a size-exclusion chromatography (SEC) Superdex 200 Increase 10/300 GL column in a SAXS buffer (20 mM Tris-HCl, 150 mM NaCl, 2 mM tris(2-carboxyethyl)phosphine (TCEP), 3% glycerol, pH 8). The resulting pure protein fractions were collected, concentrated, and subsequently adjusted to a concentration of 3 mg/ml, calculated using the formula C (mg/ml) = 100 / MW (kDa). The samples were measured at ESRF, BM29 BioSAXS of Grenoble (France). All experimental scattering data were corrected by subtracting the buffer scattering data and then fitted to the theoretically calculated curve of AlphaFold2 predicted models^23,25^ using the *FoXS* online server to calculate the quality of fit, chi² values^36^.

### X-Ray crystal structure

To improve the quality of the protein crystals by slightly decreasing the protein solubility, we redesigned the 6H5L surface by replacing 15 lysine residues on the equivalent helical wheel *f* position with glutamine (6H5L_FKQ variant), resulting in a decreased pI from 9 to 5.7^37,38^. The prediction of the 6H5L_FKQ variant was carried out using Alphafold2 and compared to the original 6H5L. 6H5L_FKQ variant was successfully produced in *E. coli* BL21 Star (DE3) competent cells and thoroughly characterized using our standard methods (Suppl Table S6).

Crystallization drops were set up with commercial crystallization screens using the vapor diffusion method employing a mosquito® Xtal3 crystallization robot (SPT Labtech) and incubated at 293 K. The protein concentration was 20 mg/ml in 20 mM Tris-HCl, 150 mM NaCl, 2 mM TCEP pH 8.0 and the drop volume was 400nL, with a 1:1.5 ratio of protein and precipitant solution. Crystallization drops were equilibrated against a reservoir containing 40 µL of precipitant solution.

The crystals were obtained from manually set up drops of 1 µL protein mixed with 1.5 µL precipitant solution (17.5% PEG 3350, pH = 7.8, 150 mM L-Malic Acid. 80 µL of precipitant solution in the reservoir). Crystals appeared after one day to two weeks. The obtained crystals were harvested from mother liquor with CryoLoops (Hampton Research) and briefly incubated with mother liquor containing 25% glycerol or 25% PEG400 followed by flash freezing in liquid nitrogen. Diffraction data was collected at 100 K on beamlines MASSIF-3 and ID30B at ESRF, Grenoble (France). Full datasets (360°) were collected to 2.8 Å resolution.

The collected data were processed using XDS^39^. with the provided input file from the beamline. Structure determination was performed by molecular replacement using PHASER^40^ with the design models as search templates. The best solution was refined in reciprocal space with PHENIX^41^ with 5% of the data used for R_free_ and by real-space fitting steps against σA-weighted 2F_o_–F_c_ and F_o_–F_c_ electron density maps using COOT^42^. Water molecules were placed automatically into difference electron density maps and accepted or rejected according to geometry criteria and their B-factors, defined in the refinement protocol. Details of data collection, processing, and refinement are summarized in Supplementary Table S7. The final coordinates of the refined model have been deposited to the wwPDB (accession code 9S1T).

### Inhibition reaction

The diketone inhibition reactions with 5 (Figure 3B) for all protein variants were carried out at 29 °C in a buffer solution (50 mM sodium phosphate, 150 mM NaCl, and 5% DMSO) at pH 8. The reaction mixture consisted of a protein concentration of 28 µM and 28-280 µM diketone **5** (1-(6- methoxy-2-naphthalenyl)-1,3-butanedione) (1:1-10 ratio) in a total reaction volume of 1 mL. After an overnight incubation period, the reaction mixture was loaded over SEC to remove the excess diketone, collecting only the protein fraction to measure. For a clear comparison, UV-Vis spectra were obtained for the free protein as well as the labeled protein reaction product.

### Competition assay

We have implemented two distinct competition assays. The first assay (DPH_diketone competition assay) involved carrying out a DPH fluorescent binding experiment, where we compared the fluorescence intensities between the free protein and the proteins labeled with diketone inhibitor. To ensure consistent protein concentrations, we standardized all the SEC fractions of the treated proteins to possess identical absorbance values at 280nm before initiating the fluorescence measurements (Figure. 4A & 5A).

The second experiment included a titration assay, in which the diketone **5** from DMSO stock solution was (up to 1.72 mM) to the saturated protein-DPH complex (1 µM of each after 1 hour of incubation with shaking at 300 rpm), and the corresponding fluorescent signals for DPH dye (Supplementary Table S2) were observed at various titration points.

Both measurements were carried out at 29 °C in a buffer solution (50 mM sodium phosphate, 150 mM NaCl, and 5% DMSO) at pH 8 using 96-Well Black Flat Bottom Polystyrene Non-Binding Surface Microplates (Corning, Lowell, MA) using a BMG Labtech instrument (Aylesbury, UK) Clariostar plate reader. To ensure accuracy and account for background effects, fluorescence signals were measured for both the buffer-DPH and buffer-DPH-diketone mixtures under identical conditions.

### Retro-aldolase activity fluorescence assay

Kinetic assays of all protein variants were conducted in 96-Well Black Flat Bottom Polystyrene Non-Binding Surface Microplates (Corning, Lowell, MA) using a plate reader (CLARIOstar, BMG Labtech). Initial reaction rates for the conversion of 4-hydroxy-4-(6-methoxy-2-naphthyl)-2- butanone **3** (methodol) to 6-methoxy-2-naphthaldehyde **1** (Figure 3A) were determined by monitoring the formation of the fluorescent product **1** with an excitation wavelength *λex* 330 nm and emission wavelength *λem* 452 nm measured from the top (with a bandwidth of 2 nm)^26^. All measurements were conducted at 29 °C in assay buffer (50 mM sodium phosphate, 150 mM NaCl, and 7.5% DMSO) at pH 8. The stock of **3** was dissolved in DMSO and stored at -20 °C. The fluorescence signal was corrected for the buffer-catalyzed background reaction under the same conditions and converted to product concentration using a calibration curve within the product formation range. The data were subjected to Michaelis–Menten fitting using GraphPad Prism 8 to determine catalytic rate constant (k_cat_) and Michaelis constant (K_M_) values.

### Aldol/retro-aldol reactions chromatographic assay

**General method:** we used the same buffer system (50 mM sodium phosphate, 150 mM NaCl, pH 8) and 500 µL total reaction volume. for both reversible reactions. The reaction conditions were kept constant at 30 °C, shaking at 120 rpm for 24 hours for all aldol and retro-aldol reactions. However, protein, substrate, and cosolvent concentrations varied depending on the experiment’s objective. The products were extracted using ethyl acetate (2 x 250 µL) spiked with 0.5 vol% acetophenone as an internal standard (IS). The combined extracts were dried over anhydrous sodium sulfate and analyzed by HPLC with a 5 µL injection volume. A Chiralpak® AS-H column (250 x 4.6 mm, 5 µm, Daicel chiral technologies) was used for activity analysis, and a Chiralcel® OD-H column (250 x 4.6 mm, 5 µm, Daicel chiral technologies) for chirality. Compound identity was confirmed by GC/MS. The results were corrected for buffer-catalyzed background reactions under the same conditions.

**Aldol reaction:** To perform the aldol reaction, 5 mM of 6-methoxy-2-naphthaldehyde **1** was added to the reaction buffer from a stock solution in acetonitrile (20 vol% final concentration as co-solvent), along with 10 vol% acetone **2** to drive the reaction forward. The reaction was carried out with a protein concentration of 100 µM.

**Retro-aldol reaction:** For the retro-aldol reaction, we used a protein concentration of 10 µM to catalyze the reaction with 1 mM Mehtodol **3** as the substrate, added from a stock solution in acetonitrile (5 vol% final concentration as co-solvent) in the reaction buffer.

**Tune the two-step reaction and product profile,** we reduced the acetonitrile concentration as a co-solvent from 20% to 1% and acetone from 10% to 5% in the aldol reaction. using the same reaction conditions, buffer system, and 100 µM protein concentrations as described above.

**Enzyme stability study:** The aldol reaction was performed with a 10 µM protein concentration, using both freshly prepared protein and protein stored for three months at 4 °C. The substrates, 5 mM 6-methoxy-2-naphthaldehyde **1**, was added from a stock solution in acetonitrile (10 vol% final concentration as co-solvent) with 5 vol% acetone **2**, under the same general reaction condition.

### Whole cells biotransformation

*E. coli* BL21 Star (DE3) cells were lyophilized after overexpressing either 6H5L or 6H5L_Top and their catalytic activity was tested in aldol and retro-aldol reactions. Empty plasmid cells were used as control. The reaction setup was as follows: 15 mg of lyophilized cells were resuspended in 300 µL reaction buffer (50 mM sodium phosphate, 150 mM NaCl, pH 7.5) for 20 min at 30 °C under shaking at 120 rpm. Subsequently, 5 mM of substrate was added to each reaction. In the aldol reaction, 6-methoxy-2-naphthaldehyde **1** was added from a stock solution in acetonitrile (10 vol% final concentration co-solvent) followed by 5 vol% acetone. In the retro-aldol reaction, 4- hydroxy-4-(6-methoxy-2-naphthyl)-2-butanone **3** (methodol) was added from a stock solution in acetonitrile (20 vol% final concentration co-solvent). The reactions (final volume 500 µL) were incubated overnight at 30 °C and 120 rpm. Control reactions were also run in the absence of cells. The products were extracted using ethyl acetate (2 x 250 µL) spiked with 0.5 vol% acetophenone as internal standard (IS). The combined extracts were dried over anhydrous sodium sulfate and analyzed by HPLC using an injection volume of 5 µL and a Chiralpak® AS-H column (250 x 4.6 mm, 5 µM, Daicel chiral technologies) for the aldol reaction and a Chiralcel® OD-H column (250 x 4.6 mm, 5 µM, Daicel chiral technologies) for the retro-aldol reaction. The identity of the compounds was further confirmed by GC/MS and the use of authentic reference materials. The results were obtained by using a calibration curve generated with authentic reference material under same conditions and corrected for the buffer-catalyzed background reaction under the same conditions.

## ACKNOWLEDGEMENTS

We thank David Baker for detailed discussion in the beginning of this project and are grateful for his sustained advice throughout. We also want to thank Dek Woolfson for inspiring conversations about surface residue compositions in coiled coils and their crystallizability. Additionally, we thank Peter Macheroux for important remarks about the kinetics of the designed system and Lukas Schober for for technical support in the synthesis of methodol. We acknowledge the European Synchrotron Radiation Facility for provision of synchrotron radiation facilities, and we would like to thank the staff of the ESRF and EMBL Grenoble for assistance and support in using beamlines MASSIF-3, ID30B, and BM29. M.H thanks the University of Graz for funding. W.E. and G.O. were supported by the Austrian Science Fund (FWF) grant 10.55776/P30826 and by funding from the European Research Council through an ERC Starting Grant (HelixMold 802217). H.L. and G.O. were supported by a FETOPEN project (ARTIBLED, 863170). This research was funded in whole, or in part, by the Austrian Science Fund (FWF) [10.55776/P30826 to GO].

## Author contributions

Conceptualization: W.E., G.O.; methodology: W.E, M.H, G.O.; Validation: W.E., B.G., D.S., H.L., M.H., G.O.; Formal analysis: W.E., M.H., G.O.; Investigation: W.E, B.G., D.S., M.C., H.L., M.A., M.H., G.O.; Resources: M.H., G.O.; Data Curation: W.E., G.O.; Writing: W.E., G.O.; Visualization: W.E., G.O; Supervision: M.H., G.O.; Project administration: G.O.; Funding acquisition: M.H., G.O.

## Data and materials availability

Supplementary Materials are available for this paper. Final design models and corresponding data have been deposited to Zenodo (DOI <u>10.5281/zenodo.16327056</u>). Correspondence and requests for materials should be addressed to G.O.

