## Supplementary Materials for "Computational design of a thermostable α-helical barrel protein with tunable active-site positioning"

**for**

**Content:**

**Protein production, characterization and activity assays**

- Supplementary Figures S1 – S12
- Supplementary Tables S1 – S7

**RosettaScripts XML[1] and Geometric Constraint File Format for RosettaMatch[2]**

**Synthesis, HPLC, GC-MS analysis and stander curves.**

#### Protein production, characterization and activity assays

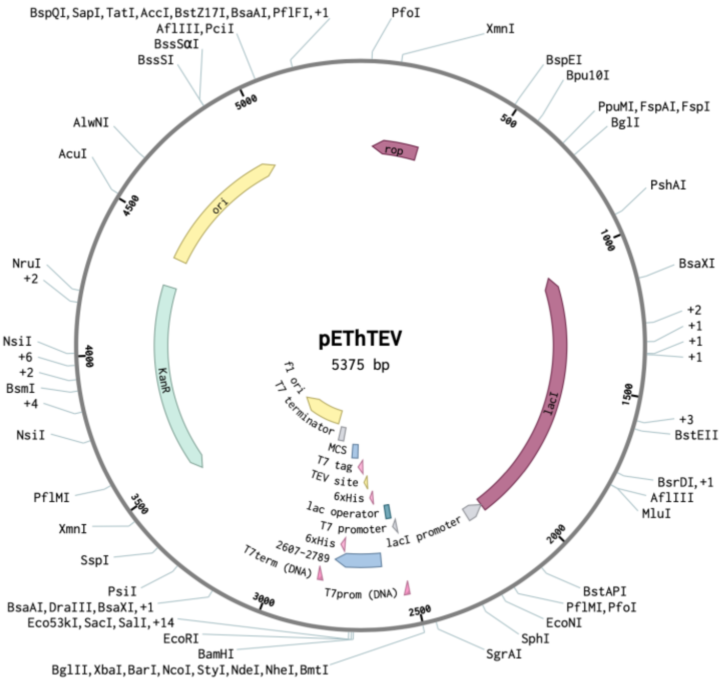


**Figure S1.** pEThTEV vector.

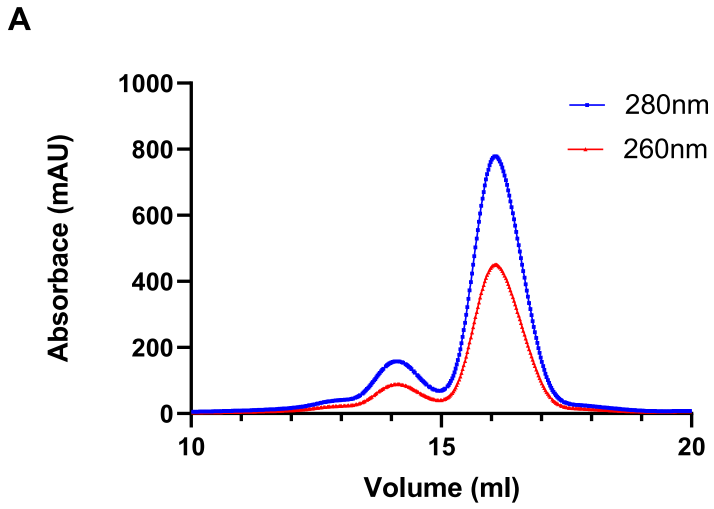


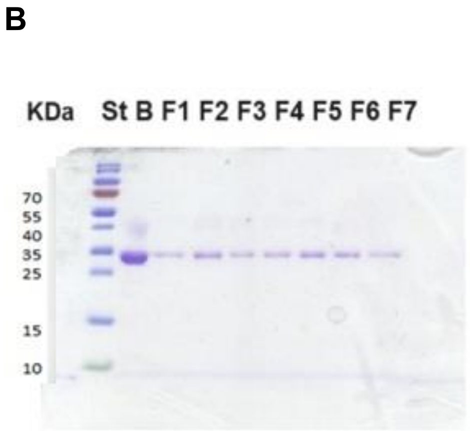


**Figure S2. A)** Size exclusion chromatogram of 6H5L desired monomer fraction as the highest peak at the expected retention volume with good separation from the dimer form. **B)** SDS-PAGE gel (St) protein ladder prestained dual color protein molecular weight marker (10-170 kDa) , (B) protein band before loading and (F1-F7) pure protein fractions from SEC.

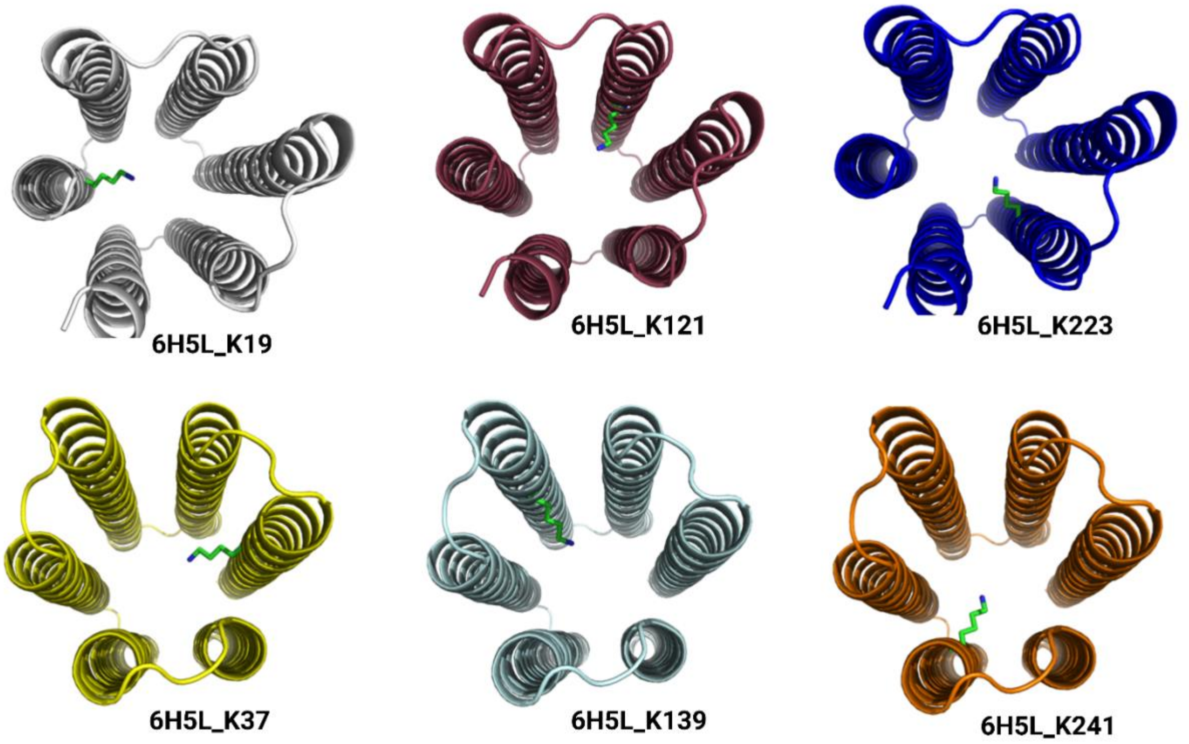


**Figure S3.** Structural representation of six single lysine mutants, each with one lysine residue introduced into the hydrophobic channel.

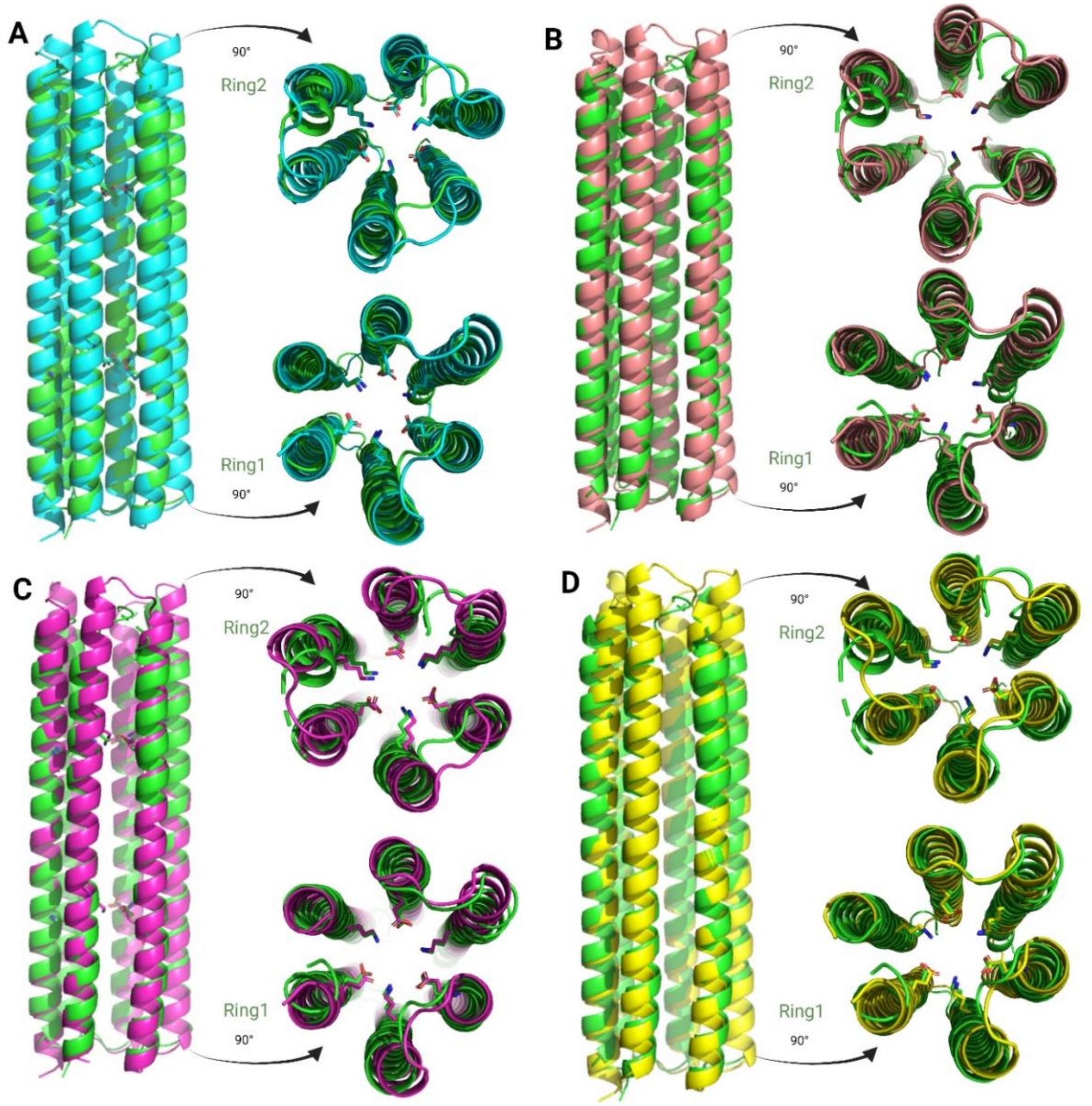


**Figure S4.** Structure representations in cartoon form (side and top view) of 6H5L_FKQ crystal structure (always in green) compared with different ML prediction methods **A)** Alphafold2 [3,4], **B)** ESMFold [3,5],**C)** OmigaFold [3,6] and **D)** RoseTTAFold [3,7] demonstrating high similarity in overall structural variations and two rings within the hydrophobic channel across predictive models and the experimental structure.

**Table S1:** Comparison of overall structure variances and two rings in the hydrophobic channel between the 6H5L_FKQ experimental structure and versus ML prediction methods in RMSD

| **Prediction method** | RMSD | | |
|---|---|---|---|
| | **Overall** | **Ring 1** | **Ring 2** |
| **AlphaFold2**[3,4] | 0.625 | 0.213 | 0.414 |
| **ESMFold**[3,5] | 0.700 | 0.350 | 0.298 |
| **OmegaFold**[3,6] | 1.029 | 0.407 | 0.692 |
| **RoseTTAFold2**[3,7] | 1.035 | 0.674 | 0.579 |

**Table S2.** Structure representation, excitation (λex) and emission (λem) wavelengths of DPH and TMA-DPH fluorescence dyes utilized for the ligand binding assay[8].

| Ligand | DPH | TMA-DPH |
|---|---|---|
| **Structure** | 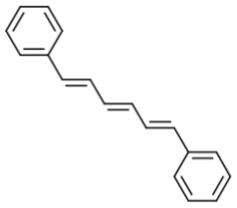 | 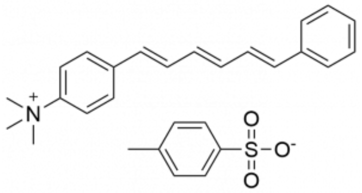 |
| **λex (nm)** | 350 ± 16 | 356 ± 16 |
| **λem (nm)** | 380 – 602 | 390 – 603 |

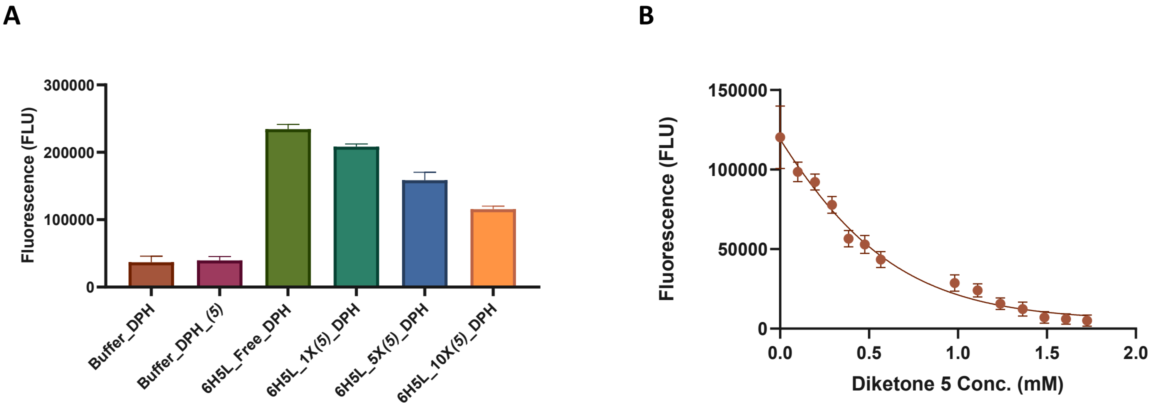


**Figure S5.** Competitive assays of DPH dye and inhibitor **5** with 6H5L design. **A)** DPH with **5** competitive assay using same protein concentration of free and different diketone **5** labeled 6H5L protein incubating with 1 μM DPH dye showing the highest fluorescence of the free 6H5L protein and fluorescence decreasing by more diketone **5** concentration (1X, 5X & 10X). **B)** Titration curve of a diketone inhibitor **5** on 6H5L protein bonded with DPH dye (both 1μM), shows a decrease in fluorescence during titration as the diketone concentration increases, with an inhibition constant (Ki) value of 0.27 mM. Both measurements were carried out at 29 °C in a buffer solution (50 mM sodium phosphate, 150 mM NaCl, and 5% DMSO) at pH 8.

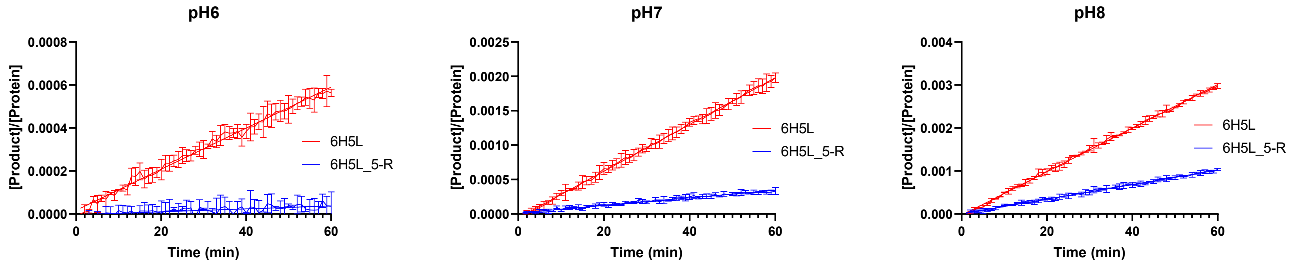


| Protein | Reaction velocity (V) (μM/min) *10^-5 | | |
|---|---|---|---|
| | **pH 6** | **pH 7** | **pH 8** |
| **6H5L** | 0.96 ±0.02 | 3.4 ±0.02 | 5.1 ±0.02 |
| **6H5L_6-R (knockout)** | 0.07 ±0.01 | 0.5 ±0.01 | 1.7 ±0.01 |

**Figure S6.** Retro-aldolase progress curves and reaction velocity of 6H5L design and its knockout version 6H5L_6-R (without lysines in the core), using the same protein concentration of 10 μM and 250 μM methanol **3** substrate at different pH levels at 29 °C in assay buffer (50 mM sodium phosphate, 150 mM NaCl, and 7.5% DMSO).

**Table S3.** Retro-aldolase kinetic parameters of all 6H5L single lysine variants.

| Protein variant | $K_M$ (µM) | $k_{cat}$ (x $10^{-4}$ $min^{-1}$) | $k_{cat}/K_M$ ($M^{-1}$ $min^{-1}$) |
|---|---|---|---|
| 6H5L_K19 | 1753 ±818 | 9.884 ±0.31 | 0.56 |
| 6H5L_K 37 | 2100 ±437 | 17.78 ±2.6 | 0.85 |
| 6H5L_K 121 | 1627 ±217 | 11.63 ±1.0 | 0.71 |
| 6H5L_K 139 | 2153 ±1150 | 15.32 ±5.8 | 0.71 |
| 6H5L_K 223 | 1307 ±494 | 9.799 ±2.3 | 0.75 |
| 6H5L_K 241 | 1015 ±240 | 8.994 ±1.2 | 0.89 |

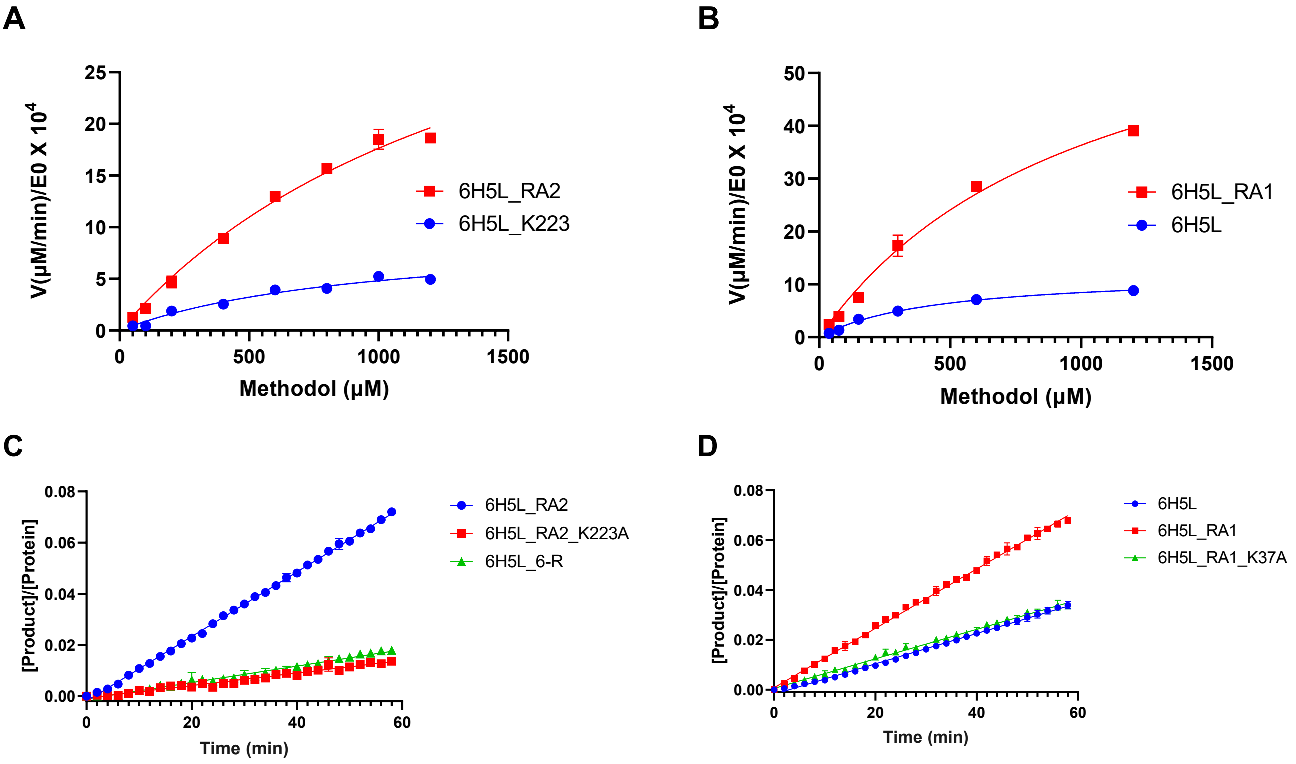


**Figure S7.** Retro-aldolase Michaelis–Menten plots and progress curves with methodol **3** as a substrate of 6H5L active site redesigned variants compared to the initial activity of the input designs. E0, total enzyme concentration; V, initial reaction velocity. **A**) 6H5L_RA2 variant: nearly 3-fold activity increase compared to 6H5L_K223. **B)** 6H5L_RA1 variant: 3.5-fold increase compared to 6H5L. (Table 3). Retro-aldolase progress curves of **C**) 6H5L_RA2 **D)** 6H5L_RA1, active site redesigned variants of 6H5L compared to and its initial input designs and knockout versions using the same protein concentration of 5 µM and 500 µM methanol **3.** All the reactions were performed at 29 °C in assay buffer (50 mM sodium phosphate, 150 mM NaCl, and 7.5% DMSO) at pH 8.

**Table S4.** reaction velocity of 6H5L and active site redesigned variants and its knockout versions (without lysines in the core), using the same protein concentration of 5 µM and 500 µM methodol as substrate.

| Protein | Reaction velocity (V) (μM/min) *10$^{-3}$ |
|---|---|
| 6H5L | 0.44 ±0.003 |
| 6H5L_Top | 3.02 ±0.008 |
| 6H5L_Top_K199A | 0.58 ±0.003 |
| 6H5L_RA2 | 1.26 ±0.006 |
| 6H5L_KRA2_K223A | 0.24 ±0.006 |
| 6H5L_6-R | 0.32 ±0.007 |
| 6H5L_RA1 | 1.19 ±0.008 |
| 6H5L_RA1_K37A | 0.59 ±0.006 |

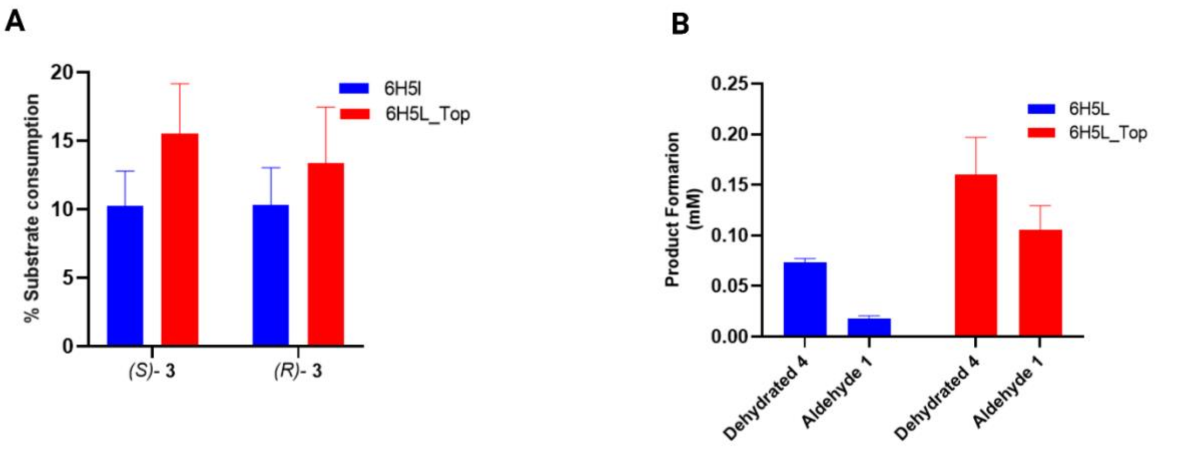


**Figure S8.** Retro-aldol reaction catalysis by 10 μM of 6H5L (blue) or 6H5L_Top (red) (50 mM sodium phosphate, 150 mM NaCl, and 5% MeCN at pH 8). **A)** Substrate consumption (%) *of (S) & (R)*-Methodol **3. B)** Product formation of 6-methoxy-2-naphthaldehyde **1** and the corresponding dehydration product **4**. Measurements were performed on chiral-phase HPLC to monitor the consumption of both enantiomers.

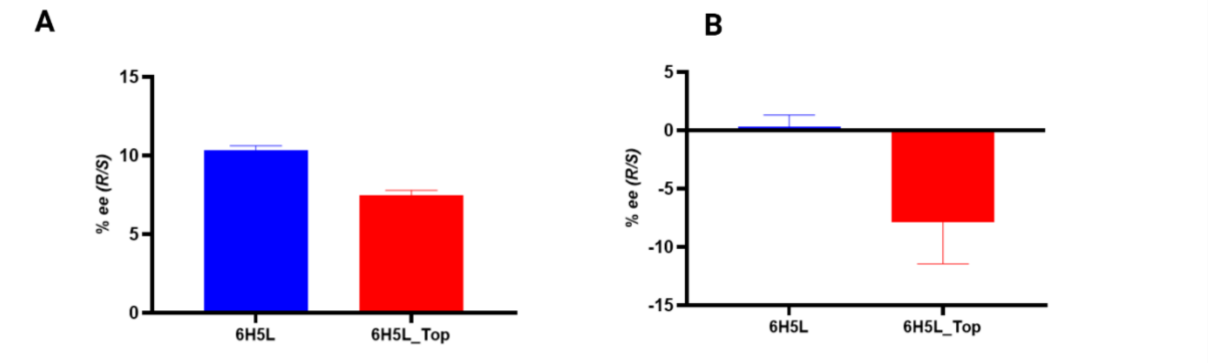


**Figure S9.** Enantiomeric excess (% ee) of *(R/S)*-Methodol **3** isomers in reactions catalyzed by 6H5L (blue) and 6H5L_Top (red). **A)** As products of the aldol reaction. **B)** As substrates in the retro-aldol reaction. Measurements were performed on chiral-phase HPLC.

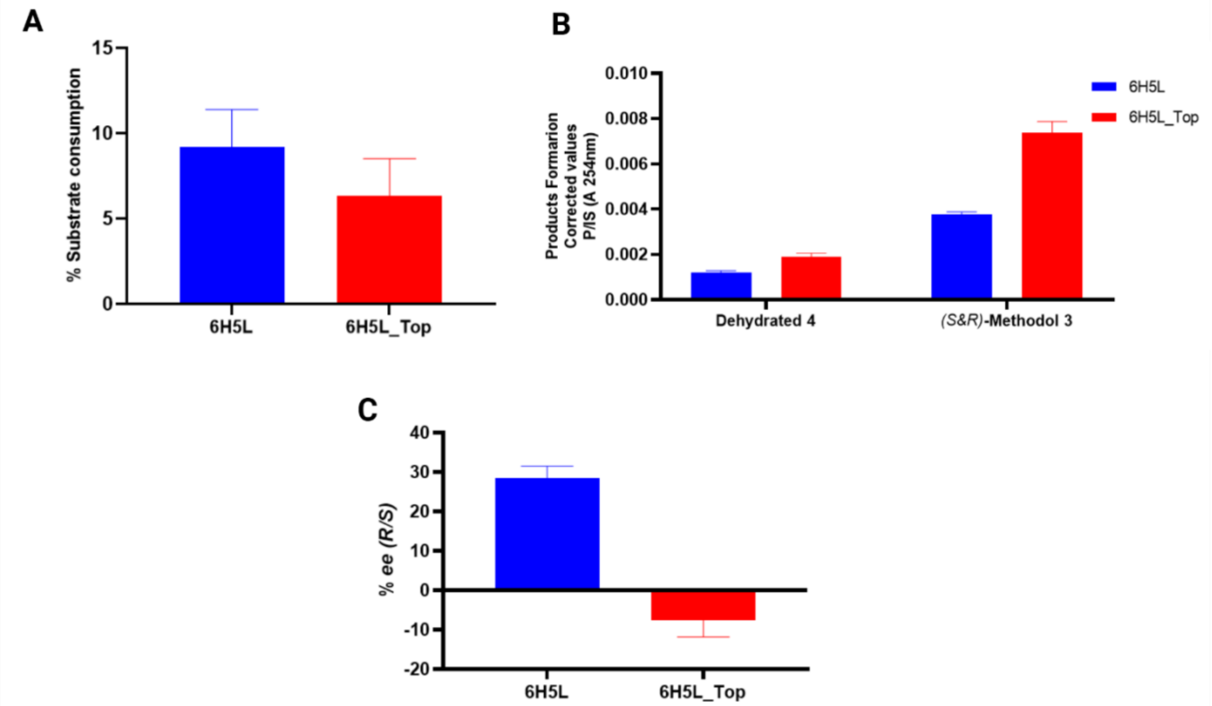


**Figure S10.** Aldol reaction catalysis by 100 µM of 6H5L (blue) or 6H5L_Top (red) using less co-solvent (1% MeCN) in buffer (50 mM sodium phosphate, 150 mM NaCl, at pH 8). **A)** % substrate consumption of 6-methoxy-2-naphthaldehyde **1**. **B)** Product formation of *S* & *R* - Methodol **3** and the corresponding dehydrated product **4**. **C)** Enantiomeric excess (% ee) of (*R/S*)- Methodol **3** isomers as substrate. Measurements were performed on chiral-phase HPLC.

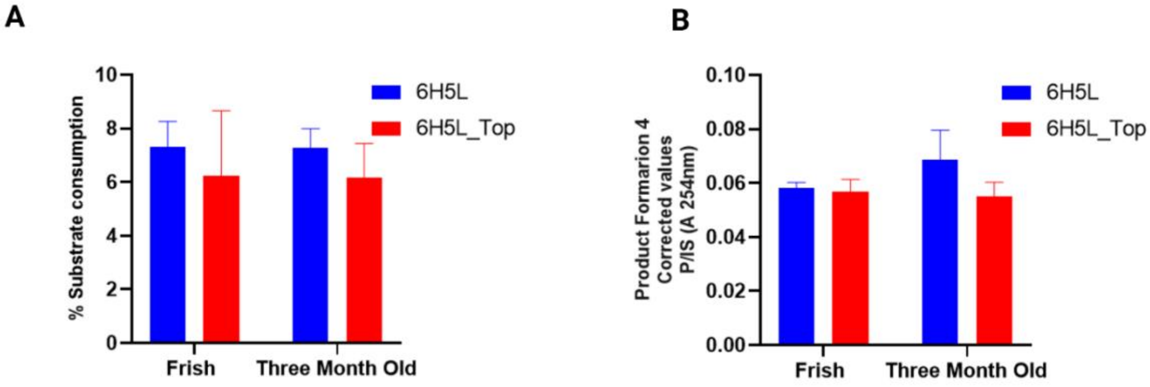


**Figure S11.** Aldol reaction catalysis by freshly produced and three-month-old 6H5L (blue) or 6H5L_Top (red) proteins (10 µM) in buffer (50 mM sodium phosphate, 150 mM NaCl, and 20% MeCN at pH 8). **A)** % substrate consumption of 6-methoxy-2-naphthaldehyde **1. B)** Formation of dehydrated product **4**. Measurements were performed on HPLC

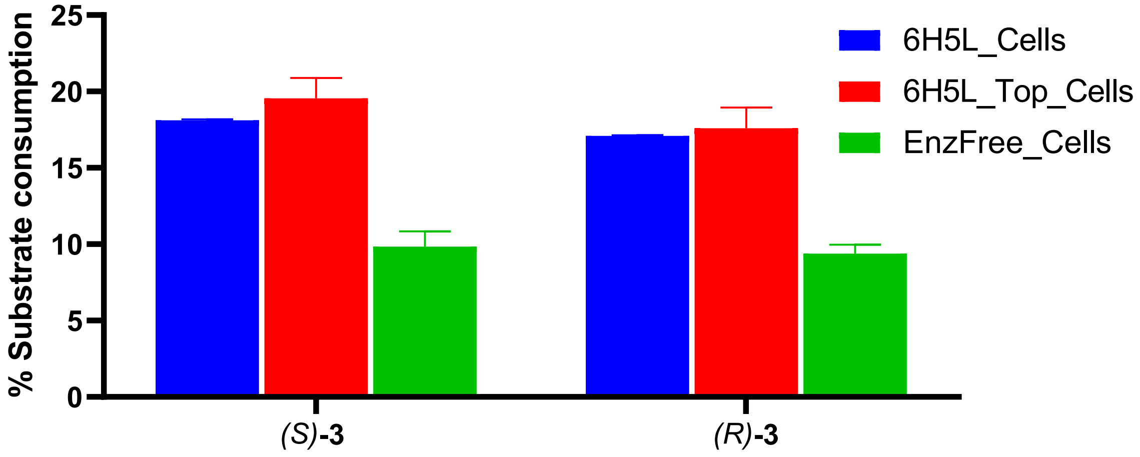


**Figure S12.** Substrate consumption in the retro-aldol reaction of 5 mM of **3** catalyzed by 6H5L and 6H5L_Top variants (whole cells) compared to enzyme free cells (EnzFree_Cells) in 50 mM sodium phosphate buffer (pH 7.5) containing 150 mM NaCl and 20 vol% acetonitrile. Measurements were performed on chiral-phase HPLC to monitor the consumption of both enantiomers.

**Table S5**. Amino acid sequences and theoretical pIs of 6H5L design and its redesign variants.

| Protein | Sequence | Theoretical pI |
| --- | --- | --- |
| **6H5L** | TCEVVKKILYMAKKLVEKKKEVVKKILKMAKELVEKKKEVVYKALEAAEKGLDTK<br>KIAKLLLEMLEHELELAEKIAKLLLEMLEEELKLAEKIAKLMLESGISEEVVKKILY<br>MAKKLVEKKKEVVKKILYMAKELVEKKKEVVKKALEAAEKGLDTKKIAKLLLEML<br>EHELELAEKIAKLLLEMLEEELKLAEKIAKLMLESGISEEVVKKILKMAKKLVEKKK<br>EVVKKILYMAKELVEKKKEVVKKALEAAEYGLDTKKIAKLLLEMLEHELELAEKIA<br>KLLLEMLEEELKLAEKIAKLMEEQCK | 9.05 |
| **6H5L_fKQ** | TCEVVKKILYMAKKLVEQKKEVVKQILKMAKELVEKKKEVVYKALEAAEKGLDT<br>KKIAKLLLEMLEHELELAEQIAKLLLEMLEEELQLAEKIAQLMLESGISEEVVKQIL<br>YMAKQLVEKKKEVVKKILYMAKELVEKKKEVVQKALEAAEKGLDTKQIAKLLLE<br>MLEHELELAEKIAQLLLEMLEEELKLAEKIAKLMLESGISEEVVKKILQMAKKLVE<br>KKKEVVQKILYMAQELVEKKQEVVKKALEAAEYGLDTKKIAQLLLEMLEHELELA<br>EKIAKLLLEMLEEELKLAEKIAKLMEEQCK | 5.72 |
| **6H5L_Top** | GCEVYKKILYMAKKLVEKKKEVVKKILKMAKELVEKKKEVVYKALEAAEKGLDT<br>KKIAKLLLEMLEHELELAEKIAKLLLEMLEEELKLAEKIAKLFLEEGAEEEVLKKWL<br>YMAKKLVEKKKEVVKKILYMAKELVEKKKEVVKKALEAAEKGLDTKKIAKLLLEM<br>LEHELELAEKIAKLLLEMLEEELKLAEKTAKLKLESGASEEIQKKYLKMAKKLVEK<br>KKEVVKKILYMAKELVEKKKEVVKKALEAAEYGLDTKKIAKLLLEMLEHELELAE<br>KIAKLLLEMLEEELKLAEKYAKLYEEQCK | 8.96 |
| **6H5L_RA1** | TCEVVKKILYMAKKLVEKKKEVVKKILKMAKELAEKKKEVVYKALEAAEKGLDTK<br>KIAKLALELLEHLLELAEKIAKLLLEMLEEELKLAEKIAKLMLESGISEEVVKKILYM<br>AKKLVEKKKEVVKKILYMAKELVEKKKEVAKKALEAAEKGLDTKKIAKLLLEMAE<br>HELELAEKIAKLLLEMLEEELKLAEKIAKLMLESGISEEVVKKILKMAKKLVEKKKE<br>VVKKILYMAKEYVEKVKEVVKKALEAAEYGLDTKKIAKLLLEYLEHQLELAEKIAK<br>LLLEMLEEELKLAEKIAKLMEEQCK | 9.10 |
| **6H5L_RA2** | TCEVVKKILYMAKKLVEKRKEVVKKILKMAKELVEKRKEVVYKALEAAEKGLDTK<br>KIAKLLLEMLEHELELAEKIAKLLLEMLEEELKLAEKIAKLMLESGISEEVVKKILY<br>MAKKLVEKRKEVVKKILYMAKELVEKRKEVVKKALEAAEKGLDTKKIAKLLLEML<br>EHELELAEKIAKLLLEMLEEYLKLAEKIAKLMLESGISEEVVKKILKMAKKYVEKK<br>KEVVKKILYMAKELVEKRKEVVKKALEAAEYGLDTKKIAKLLLEMLEHELELAEKI<br>AKLLLEMLEEQLKLAEKIAKLMEEQCK | 9.17 |

**Table S6.** Summary of **CD** spectra, temperature scans (**CD_Temp**), **DPH** binding curves with corresponding $K_D$ values, and **SAXS** fitting results (theoretical curves in red & measured data points in blue) with accompanying chi-square values, for all 6H5L design variants.

| Design | CD | CD_Temp | DPH_Assay | SAXS |
|---|---|---|---|---|
| **6H5L_K19** | 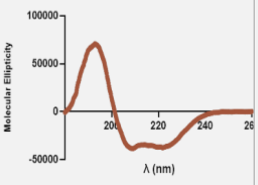  | 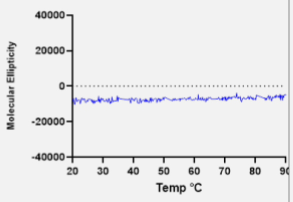  | 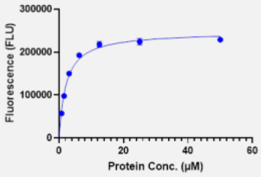  **$K_D$ = 2.16±0.13** | 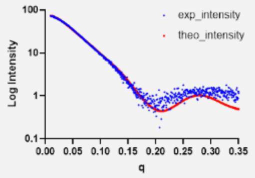  **chi² 3.0** |
| **6H5L_K37** | 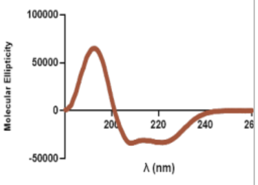  | 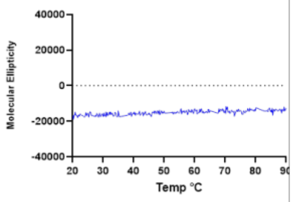  | 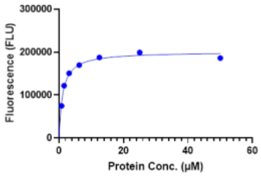  **$K_D$ = 1.12±0.07** | 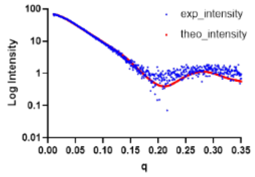  **chi² 2.2** |
| **6H5L_K121** | 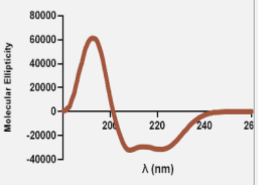  | 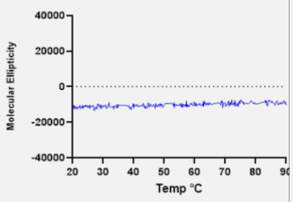  | 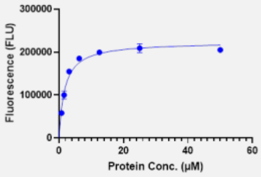  **$K_D$ = 1.72±0.14** | 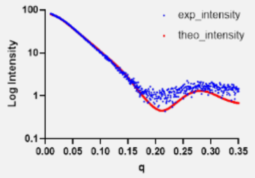  **chi² 4.1** |
| **6H5L_K139** | 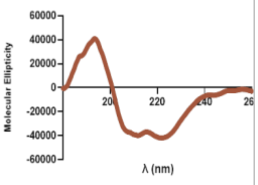  | 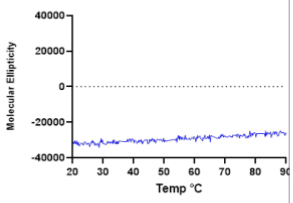  | 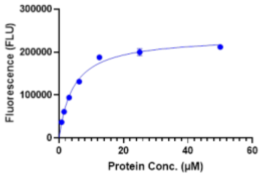  **$K_D$ = 4.45±0.29** | 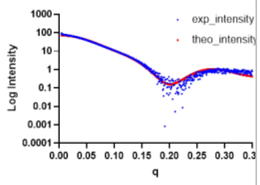  **chi² 4.6** |
| **6H5L_K223** | 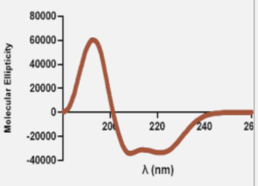  | 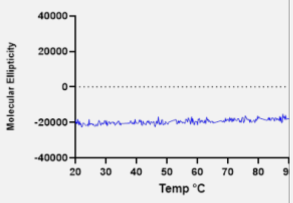  | 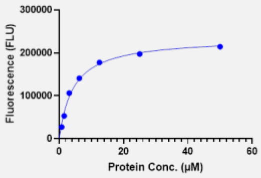  **$K_D$ = 4.36±0.24** | 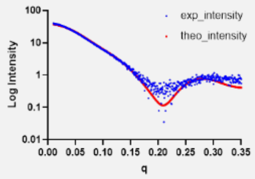  **chi² 3.5** |
| **6H5L_K241** | 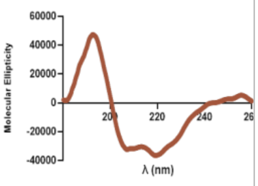  | 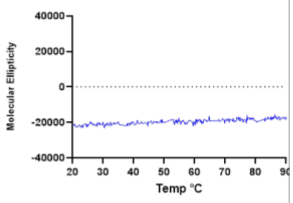  | 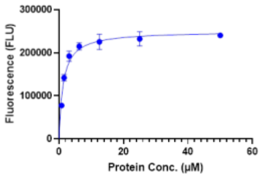  **$K_D$ = 1.26±0.13** | 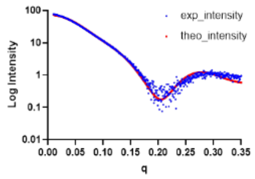  **chi² 3.6** |
| **6H5L_fKQ** | 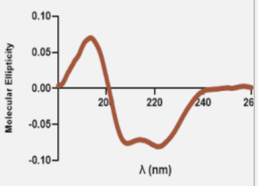  | 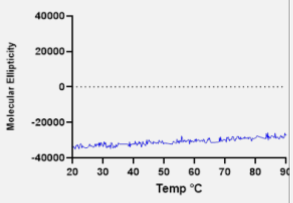  | 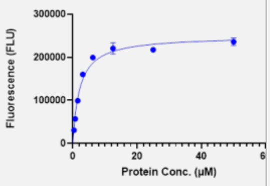  **$K_D$ = 2.11±0.16** | 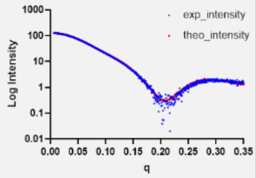  **chi² 2.2** |

| | | | | |
|---|---|---|---|---|
| **6H5L_6-R** | 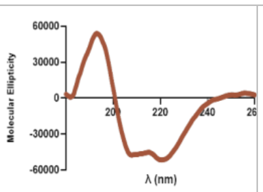 | 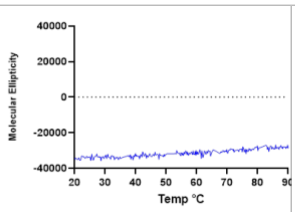 | 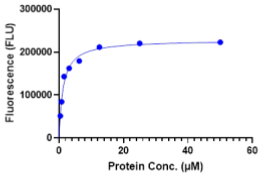 ***K*<sub></sub>D = 1.21±0.07** | 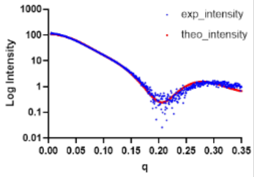 **chi²** 4.73 |
| **6H5L_RA2** | 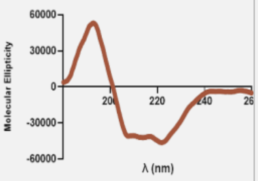 | 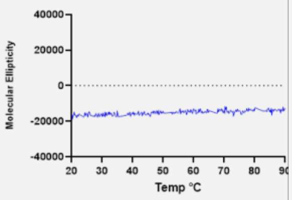 | 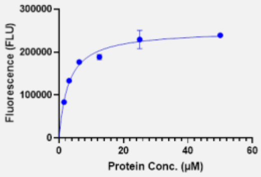 ***K*D = 2.95±0.27** | 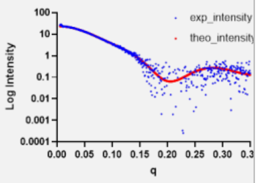 **chi²** 0.92 |
| **6H5L_RA1** | 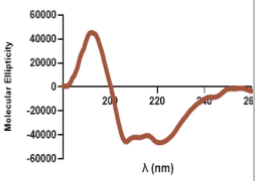 | 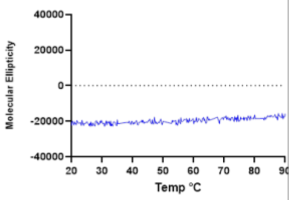 | 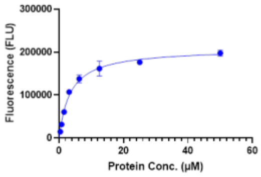 ***K*D = 3.48±0.23** | 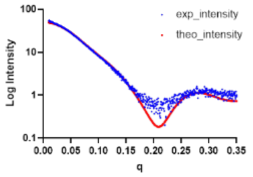 **chi²** 4.1 |
| **6H5L_Top** | 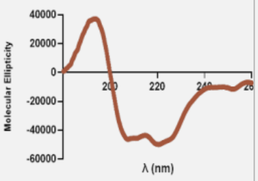 | 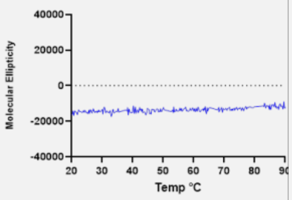 | 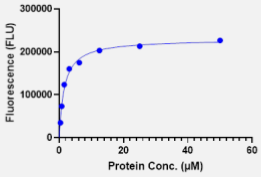 ***K*D = 1.57±0.08** | 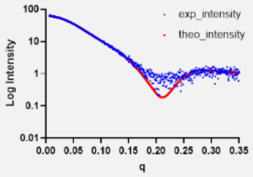 **chi²** 1.5 |

**Table S7**. Crystallographic data.

| | **6H5L_fKQ** |
|---|---|
| **Wavelength** | 0.8856 |
| **Resolution range** | 36.91 - 2.798 (3.01 - 2.8) |
| **Space group** | P 1 |
| **Unit cell** | 36.567 38.6 59.452 101.11 93.99 101.92 |
| **Total reflections** | 26278 (5222) |
| **Unique reflections** | 14388 (2910) |
| **Multiplicity** | 1.8 (1.8) |
| **Completeness (%)** | 96.33 (97.01) |
| **Mean I/sigma(I)** | 8.23 (3.59) |
| **Wilson B-factor** | 63.32 |
| **R-merge** | 0.04879 (0.1098) |
| **R-meas** | 0.069 (0.1553) |
| **R-pim** | 0.04879 (0.1098) |
| **CC1/2** | 0.994 (0.986) |
| **CC*** | 0.999 (0.996) |
| **Reflections used in refinement** | 7352 (1461) |
| **Reflections used for R-free** | 739 (146) |
| **R-work** | 0.2624 (0.2507) |
| **R-free** | 0.3188 (0.3257) |
| **Number of non-hydrogen atoms** | 2390 |
| **macromolecules** | 2320 |
| **ligands** | 41 |
| **solvent** | 29 |
| **Protein residues** | 287 |
| **RMS(bonds)** | 0.001 |
| **RMS(angles)** | 0.27 |
| **Ramachandran favored (%)** | 98.57 |
| **Ramachandran allowed (%)** | 1.43 |
| **Ramachandran outliers (%)** | 0.00 |
| **Rotamer outliers (%)** | 1.95 |
| **Clashscore** | 3.41 |
| **Average B-factor** | 75.61 |
| **macromolecules** | 75.47 |
| **ligands** | 88.31 |
| **solvent** | 68.78 |

### Rosetta Scripts xml for closing loops and sequence design

```
<ROSETTASCRIPTS>
	<SCOREFXNS>
		<ScoreFunction name="ref15" weights="ref2015" >
			<Reweight scoretype="hbnet" weight="3" />
			<Reweight scoretype="buried_unsatisfied_penalty" weight="1.0" />
			<Reweight scoretype="aa_composition" weight="1.0" />
		</ScoreFunction>
		<ScoreFunction name="sfxn_soft" weights="ref2015_soft" >
		</ScoreFunction>
		<ScoreFunction name="sfxn_cart" weights="ref2015_cart" >
		</ScoreFunction>
	</SCOREFXNS>
	<RESIDUE_SELECTORS>
		# Defines and Selects layers by certain cutoff
		<Layer name="corelayer" select_core="true" select_boundary="false" select_surface="false"
use_sidechain_neighbors="true" core_cutoff="5.2" surface_cutoff="2.0" />
    <Layer name="boundarylayer" select_core="false" select_boundary="true" select_surface="false"
use_sidechain_neighbors="true" core_cutoff="5.2" surface_cutoff="2.0"/>
    <Layer name="surfacelayer" select_core="false" select_boundary="false" select_surface="true"
use_sidechain_neighbors="true" core_cutoff="5.2" surface_cutoff="2.0" />
		<ResidueName name="Prolins" residue_name3="PRO" />
	# selects only the loops (not used so far)
		<SecondaryStructure name="loops" overlap="0" use_dssp="true" ss="L" />
	# selects everything but loops (not used so far)
		<Not name="notloop" selector="loops" />
	</RESIDUE_SELECTORS>
##########################
	<PACKER_PALETTES>
	</PACKER_PALETTES>
#########################
	<TASKOPERATIONS>
	# taskoperation that determines the AA used in the respective layer
			<OperateOnResidueSubset name="coreAA" selector="corelayer" >
				<RestrictAbsentCanonicalAASRLT aas="VALIFW" />
			</OperateOnResidueSubset>
			<OperateOnResidueSubset name="boundaryAA" selector="boundarylayer" >
				<RestrictAbsentCanonicalAASRLT aas="ADEFGHIKLNQRSTVWY" />
			</OperateOnResidueSubset>
			<OperateOnResidueSubset name="surfaceAA" selector="surfacelayer" >
				<RestrictAbsentCanonicalAASRLT aas="DEGHKNQRSTWY" />
			</OperateOnResidueSubset>
		# Turns off Design and packs only at the respective layer
			<OperateOnResidueSubset name="packcore" selector="corelayer" >
				<RestrictToRepackingRLT /> # turns off design at corelayer
			</OperateOnResidueSubset>
			<OperateOnResidueSubset name="packboundary" selector="boundarylayer" >
				<RestrictToRepackingRLT />
			</OperateOnResidueSubset>
			<OperateOnResidueSubset name="packsurface" selector="surfacelayer" >
				<RestrictToRepackingRLT />
			</OperateOnResidueSubset>
		# Helix breaker Prolins packing
			<OperateOnResidueSubset name="packPs" selector="Prolins" >
			<RestrictToRepackingRLT />
			</OperateOnResidueSubset>
		#Prevent Repacking Turns off design and repacking on respective layer
			<OperateOnResidueSubset name="nodesignnorepackcore" selector="corelayer" >
				<PreventRepackingRLT /> # turn off design and repacking on core
			</OperateOnResidueSubset>
			<OperateOnResidueSubset name="nodesignnorepackboundary" selector="boundarylayer" >
				<PreventRepackingRLT />
			</OperateOnResidueSubset>
			<OperateOnResidueSubset name="nodesignnorepacksurface" selector="surfacelayer" >
				<PreventRepackingRLT />
			</OperateOnResidueSubset>
		#Extrachi enables more Roatamers costs lots of computing power
			ExtraRotamersGeneric name="extrachi" ex1="1" ex2="1" ex1aro="1" ex2aro="1"
extrachi_cutoff="5" />
	</TASKOPERATIONS>
	<MOVE_MAP_FACTORIES>
	</MOVE_MAP_FACTORIES>
	<SIMPLE_METRICS>
		#measures the Energy per Residue in the loop regions
			<PerResidueEnergyMetric name="e_per_res" output_as_pdb_nums="true"
residue_selector="loops" scoretype="total_score" scorefxn="ref15" />
```

```
	</SIMPLE_METRICS>
	<FILTERS>
		# Filters by Energy per Residues in the loop regions
			<SimpleMetricFilter name="looprese" epsilon=".0001" metric="e_per_res" comparison_type="lt"
cutoff="1.0" composite_action="all" use_cached_data="true" cache_prefix="LOOP" fail_on_missing_cache="true"
confidence="1.0" />
	</FILTERS>
	<MOVERS>
		#soft design and repacking of c, b and s
			<PackRotamersMover name="design_soft_c" nloop="1" scorefxn="sfxn_soft"
task_operations="coreAA,nodesignnorepacksurface,nodesignnorepackboundary,packPs" /> # design on corelayer, turn off d
and rp on surface and boundary
			<PackRotamersMover name="design_soft_b" nloop="1" scorefxn="sfxn_soft"
task_operations="boundaryAA,nodesignnorepacksurface,packcore,packPs" /> # design on boundarylayer, turn off d and rp on
surface, pack core
			<PackRotamersMover name="design_soft_s" nloop="1" scorefxn="sfxn_soft"
task_operations="surfaceAA,nodesignnorepackcore,packboundary,packPs" /> #design on surfacelayerlayer, pack boundary
		#hard design and packing of c b and s
			<PackRotamersMover name="design_hard_c" nloop="1" scorefxn="ref15"
task_operations="coreAA,nodesignnorepacksurface,packboundary,packPs" /> #design on core, turn off design and repack at
boundary and surface
			<PackRotamersMover name="design_hard_b" nloop="1" scorefxn="ref15"
task_operations="boundaryAA,packcore,packsurface,packPs" />
			<PackRotamersMover name="design_hard_s" nloop="1" scorefxn="ref15"
task_operations="surfaceAA,nodesignnorepackcore,packboundary,packPs" />
		#Minimizer
			<MinMover name="min_soft_cart" scorefxn="sfxn_cart" cartesian="true" bb="0" chi="1" />
			<MinMover name="min_soft" scorefxn="sfxn_soft" cartesian="false" bb="0" chi="1" />
			<MinMover name="min_hard_cart" scorefxn="sfxn_cart" cartesian="true" bb="0" chi="1" />
			<MinMover name="min_hard" scorefxn="ref15" cartesian="false" bb="0" chi="1" />
		#Loop design
			<ConnectChainsMover name="loop_closureAB" loopLengthRange="3,5"
		RMSthreshold="3"
		resAdjustmentRangeSide1="-3,4"
		resAdjustmentRangeSide2="0,0"
		chain_connections="[A+B+C+D+E]" />
		#simple metric mover that hmm BRAUCH ICH DAS ÜBERHAUPT?
				<RunSimpleMetrics name="run_loopresenergy" metrics="e_per_res" prefix="LOOP"
override="false" />
		# Improve Sequence Composition
			<AddHelixSequenceConstraints name="helixcomp" desired_min_hydrophobic_fraction="0.10" />
			<AddCompositionConstraintMover name="aacomp" selector="notloop" >
					<Comp entry="PENALTY_DEFINITION; TYPE TRP; ABSOLUTE 1;
PENALTIES 20 0 1 20; DELTA_START -1; DELTA_END 2;  BEFORE_FUNCTION QUADRATIC; AFTER_FUNCTION
QUADRATIC; END_PENALTY_DEFINITION" />
			</AddCompositionConstraintMover>
	</MOVERS>
	<PROTOCOLS>
		#close loops
			<Add mover="loop_closureAB" />
		#aacomp helix
			<Add mover="helixcomp" />
		#design soft
			<Add mover="design_soft_c" />
			<Add mover="design_soft_b" />
			<Add mover="design_soft_s" />
		#minimize soft and hard
			<Add mover="min_soft_cart" />
			<Add mover="min_soft" />
			<Add mover="min_hard_cart" />
			<Add mover="min_hard" />
		#design hard
			<Add mover="design_hard_c" />
			<Add mover="design_hard_b" />
			<Add mover="design_hard_s" />
		#minimize soft and hard
			<Add mover="min_soft_cart" />
			<Add mover="min_soft" />
			<Add mover="min_hard_cart" />
			<Add mover="min_hard" />
			<Add mover="run_loopresenergy" />
			Add filter="looprese" />
	</PROTOCOLS>
	<OUTPUT scorefxn="ref15" />
</ROSETTASCRIPTS>
```

### Rosetta Script xml for retro aldolase active site design

```
<ROSETTASCRIPTS>

  <SCOREFXNS>
    <ScoreFunction name="score15" weights="ref2015" >
            <Reweight scoretype="atom_pair_constraint" weight="1"/>
            <Reweight scoretype="angle_constraint" weight="1"/>
            <Reweight scoretype="dihedral_constraint" weight="1"/>
    </ScoreFunction>
    <ScoreFunction name="r15_blank" weights="ref2015" />
  </SCOREFXNS>

  <RESIDUE_SELECTORS>
   Chain name="chA" chains="A"/>
   <JumpDownstream name="ligand" jump="1"/>
   <Not name="not_ligand" selector="ligand" />
   <And name="ligand_contacts" selectors="not_ligand" >
      <CloseContact residue_selector="ligand" contact_threshold="5.0" />
   True name="true" />
  </And>
  </RESIDUE_SELECTORS>
  <SIMPLE_METRICS>
      <InteractionEnergyMetric name="ligand_int_score" scorefxn="r15_blank" custom_type="lig_score"
residue_selector="ligand" residue_selector2="ligand_contacts" />
  </SIMPLE_METRICS>
  <TASKOPERATIONS>
            <DetectProteinLigandInterface name="liginterface" cut1="6" cut2="8" cut3="10" cut4="12"
design="1" design_to_cys="0" />
      <SetCatalyticResPackBehavior name="cstres" fix_catalytic_aa="1" />
      <RestrictIdentities name="restrict" identities="CYS,LYS,GLU,TYR,GLN" prevent_repacking="0" />
      <RestrictAbsentCanonicalAAS name="disallow" keep_aas="AVLINQDEG"/>
      ResfileCommandOperation name="allow" command="PIKAA AVLINQDEG" residue_selector="true" />
  </TASKOPERATIONS>

 <FILTERS>
   <ScoreType name="check_distance" score_type="atom_pair_constraint" threshold="40.0" scorefxn="score15"
confidence="0.0" />
   <ScoreType name="check_angle" score_type="angle_constraint" threshold="80.0" scorefxn="score15" confidence="0.0" />
   <ScoreType name="check_dihedral" score_type="dihedral_constraint" threshold="120.0" scorefxn="score15"
confidence="0.0" />
   <SimpleMetricFilter name="filter_ligscore" epsilon=".0001" metric="ligand_int_score" comparison_type="lt" cutoff="-18.0"
use_cached_data="false" confidence="0.0" />
 </FILTERS>

  <MOVERS>
              MutateResidue name="res12" target="12" new_res="HIS_D" preserve_atom_coords="true" mutate_self="false"
update_polymer_bond_dependent="true"/> it works alone
    <AddOrRemoveMatchCsts name="cstadd" cst_instruction="add_new" cstfile="Cst_6H5L_Diket_final.cst"/>
              <EnzRepackMinimize name="6H5L_Dike_opt" scorefxn_repack="score15" scorefxn_minimize="score15"
cst_opt="1" design="0" repack_only="0" fix_catalytic="1" minimize_rb="1" minimize_bb="1" minimize_sc="1" minimize_lig="1"
min_in_stages="1" backrub="0" cycles="5" task_operations="liginterface,cstres,restrict,disallow" />
              <EnzRepackMinimize name="6H5L_Dike_des" scorefxn_repack="score15" scorefxn_minimize="score15"
cst_opt="0" design="1" repack_only="0" fix_catalytic="1" minimize_rb="0" minimize_bb="1" minimize_sc="1" minimize_lig="1"
min_in_stages="1" backrub="0" cycles="5" task_operations="liginterface,cstres,restrict,disallow" />

  </MOVERS>

  <APPLY_TO_POSE>
  </APPLY_TO_POSE>

  <PROTOCOLS>
              Add mover="res12"/>
              <Add mover="cstadd"/>
              <Add mover="6H5L_Dike_opt"/>
              <Add mover="6H5L_Dike_des"/>
              <Add filter="check_distance" />
              <Add filter="check_angle" />
              <Add filter="check_dihedral" />
              <Add filter="filter_ligscore" />

  </PROTOCOLS>

 </ROSETTASCRIPTS>
```

#### Geometric Constraint File Format (.cst File) for Matching and Enzyme Design

```
CST::BEGIN
  TEMPLATE::   ATOM_MAP: 1 atom_name: C13 C12 C11
  TEMPLATE::   ATOM_MAP: 1 residue3: SRO

  TEMPLATE::   ATOM_MAP: 2 atom_name: NZ CE CD
  TEMPLATE::   ATOM_MAP: 2 residue1: K

  CONSTRAINT:: distanceAB:    1.50   0.20 500.00  1       2
  CONSTRAINT::    angle_A:  107.30  10.00 100.00  360.00   1
  CONSTRAINT::    angle_B:  152.50  10.00 100.00  360.00   1
  CONSTRAINT::  torsion_A:  -87.70   5.00 100.00  360.00   1
  CONSTRAINT::  torsion_B: -148.70  10.00 100.00  360.00   1
  CONSTRAINT:: torsion_AB: -110.20  10.00 100.00  360.00   1
CST::END

CST::BEGIN
  TEMPLATE::   ATOM_MAP: 1 atom_name: O1 C11 C12
  TEMPLATE::   ATOM_MAP: 1 residue3: SRO

  TEMPLATE::   ATOM_MAP: 2 atom_name: OH CZ CE1
  TEMPLATE::   ATOM_MAP: 2 residue1: Y

  CONSTRAINT:: distanceAB:    1.00   0.10 500.00   1      2
  CONSTRAINT::    angle_A:  128.70  20.00 100.00  120.00   1
  CONSTRAINT::    angle_B:   93.30  10.00 100.00  120.00   1
  CONSTRAINT::  torsion_A:  -82.80  10.00 100.00  120.00   1
  CONSTRAINT::  torsion_B:  101.40  10.00 100.00  120.00   1
  CONSTRAINT:: torsion_AB: -165.90  20.00 100.00  120.00   1
CST::END

CST::BEGIN
  TEMPLATE::   ATOM_MAP: 1 atom_name: C13 C12 C11
  TEMPLATE::   ATOM_MAP: 1 residue3: SRO

  TEMPLATE::   ATOM_MAP: 2 atom_name: OH CZ CE1
  TEMPLATE::   ATOM_MAP: 2 residue1: Y

  CONSTRAINT:: distanceAB:    3.80   0.20 500.00  0       2
  CONSTRAINT::    angle_A:   83.90  10.00 100.00  360.00   1
  CONSTRAINT::    angle_B:  158.60  10.00 100.00  360.00   1
  CONSTRAINT::  torsion_A:   57.20   5.00 100.00  360.00   1
  CONSTRAINT::  torsion_B: -156.30  10.00 100.00  360.00   1
  CONSTRAINT:: torsion_AB:   73.60  10.00 100.00  360.00   1
CST::END

CST::BEGIN
  TEMPLATE::   ATOM_MAP: 1 atom_name: C13 C12 C11
  TEMPLATE::   ATOM_MAP: 1 residue3: SRO

  TEMPLATE::   ATOM_MAP: 2 atom_name: OE1 CD NE2
  TEMPLATE::   ATOM_MAP: 2 residue1: Q

  CONSTRAINT:: distanceAB:    2.70   0.20 500.00  0       2
  CONSTRAINT::    angle_A:  115.30  10.00 100.00  360.00   1
  CONSTRAINT::    angle_B:  134.10  10.00 100.00  360.00   1
  CONSTRAINT::  torsion_A:  17.60    5.00 100.00  360.00   1
  CONSTRAINT::  torsion_B:  69.40    5.00 100.00  360.00   1
  CONSTRAINT:: torsion_AB: -35.90    5.00 100.00  360.00   1
CST::END
```

#### Synthesis, HPLC and GC-MS Analysis

#### Synthesis

NMR spectra were measured on a Bruker Avance III 300 MHz NMR spectrometer. Chemical shifts are reported in ppm relative to TMS (δ = 0.00 ppm) and the coupling constants (J) in Hertz (Hz).

##### ***rac*-4-Hydroxy-4-(6-methoxy-2-naphthalenyl)-2-butanone (methodol) 3**[9]

6-Methoxy-2-naphthaldehyde **1** (500 mg, 2.69 mmol, 1 eq) was added to a 1:4 mixture of acetone (**2**, 5.4 mL) and aqueous phosphate solution (22 mL, 10 mM $NaH_2PO_4$, 111 mM NaCl, 2.7 mM KCl in water, pH 7.4) (1:4). L-Proline (62.2 mg, 0.2 eq.) was added to the solution and the reaction was stirred at room temperature for 48 h. Since the reaction had not proceeded much based on control by TLC, after 48 h, more acetone (5.4 mL) and L-Proline (62.2 mg) were added and the reaction was kept stirring for another 48 h, after which conversion appeared complete on TLC. Purification by flash chromatography (cyclohexane/ethyl acetate, 2:1) and evaporation of the solvent under reduced pressure yielded the final product **3** as a white solid (400 mg, 1.64 mmol) in 61% yield. The spectral data were in accordance with the literature.[10] **$^1$H NMR (300 MHz, $CDCl_3$)** δ 7.80 – 7.69 (m, 3H), 7.44 (dd, *J* = 8.4, 1.8 Hz, 1H), 7.16 (m, 2H), 5.31 (dd, *J* = 8.7, 3.7 Hz, 1H), 3.94 (s, 3H), 3.28 (s, 1H), 3.06 – 2.83 (m, 2H), 2.23 (s, 3H).

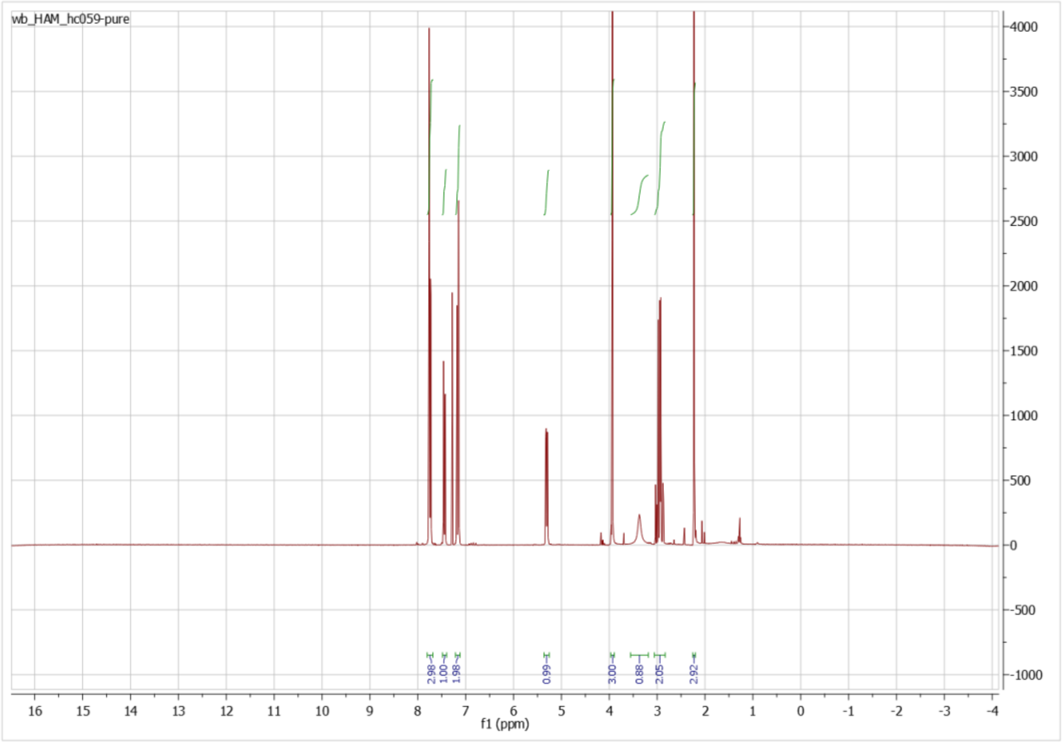


#### HPLC analysis

HPLC analyses were performed on a Shimadzu system (DGU-20A On-line Degasser, LC-20AD pump, SIL-20AC autosampler, CBM-20A system controller, SPD-M20A Photodiode Array Detector, Shimadzu CTO-20AC Column Oven). The samples (5 µL) were analyzed with an isocratic flow using heptane/*i*-propanol 92:8 as eluent at 1 mL/min for 30 min at 30 °C. Analysis was carried out at 200, 231 or 254 nm.

#### HPLC traces from representative samples

For most reactions, analysis was performed using (Daicel Chiralcel OD-H (250 mm, ID 4.6 mm, particle size 5 µm)), retention times were as follow: IS 4.98 min, 6-methoxy-2-naphthaldehyde **1** 7.52 min, dehydrated product **4** 10.87 min, 4-hydroxy-4-(6-methoxy-2-naphthalenyl)-2-butanone (*R*/*S*)-**3** 17.97 min, 4-hydroxy-4-(6-methoxy-2-naphthalenyl)-2-butanone (*S*/*R*)-**3** 19.23 min.

For aldol reactions conducted with whole-cell biotransformations, analysis was carried out using (Daicel Chiralpak AS-H (250 mm, ID 4.6 mm, particle size 5 µm)), retention times were as follow: IS 5.82 min, 6-methoxy-2-naphthaldehyde **1** 15.20 min, 4-hydroxy-4-(6-methoxy-2-naphthalenyl)-2-butanone **3** / dehydrated product **4** 19.77 min. The dehydrated product **4** and **3** co-eluted and could not be separated using this method.

#### Enzymatic catalysis by purified enzyme

Reference *rac*-4-hydroxy-4-(6-methoxy-2-naphthalenyl)-2-butanone *rac*-**3**

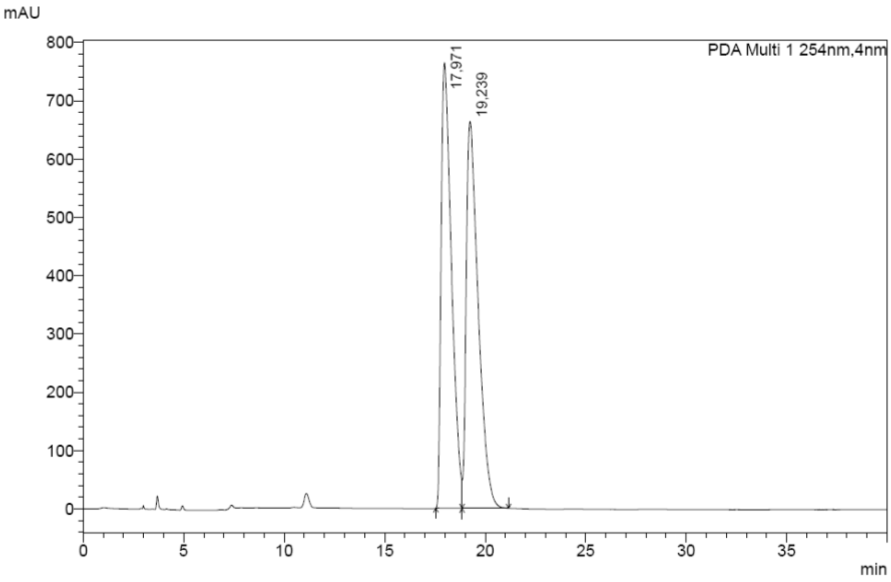


Reference dehydrated derivative 4

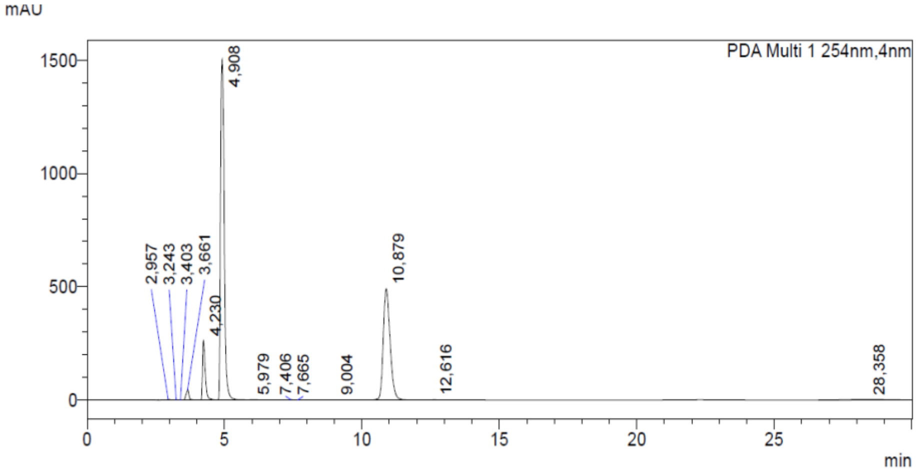

**Aldol reaction**

Retention times were as follow: IS 4.9 min, **1** 7.5 min, **4** 10.8 min, (*R*/*S*)-**3** 17.9 min, (*S*/*R*)-**3** 19.1 min.

Sample from control reaction (Blank)

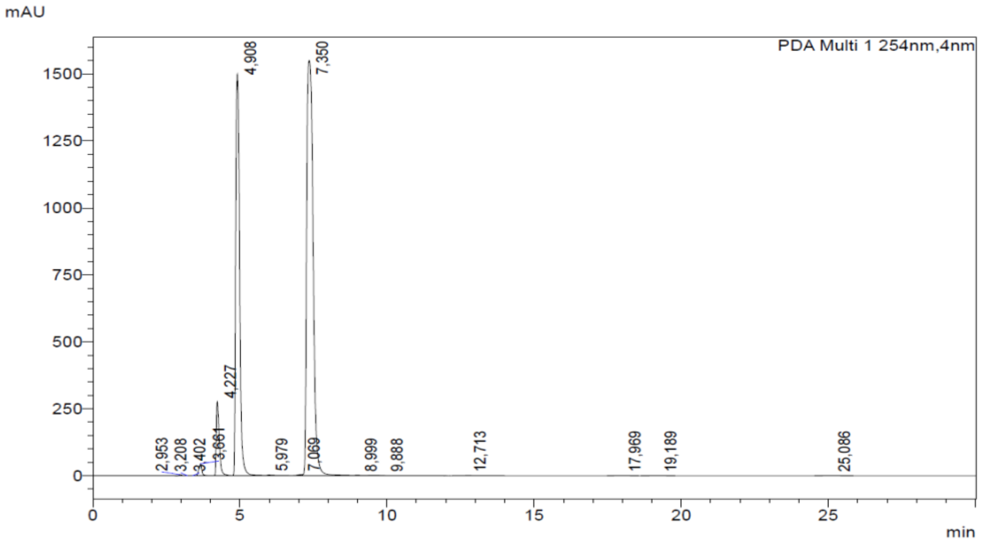


Sample from 6H5L protein

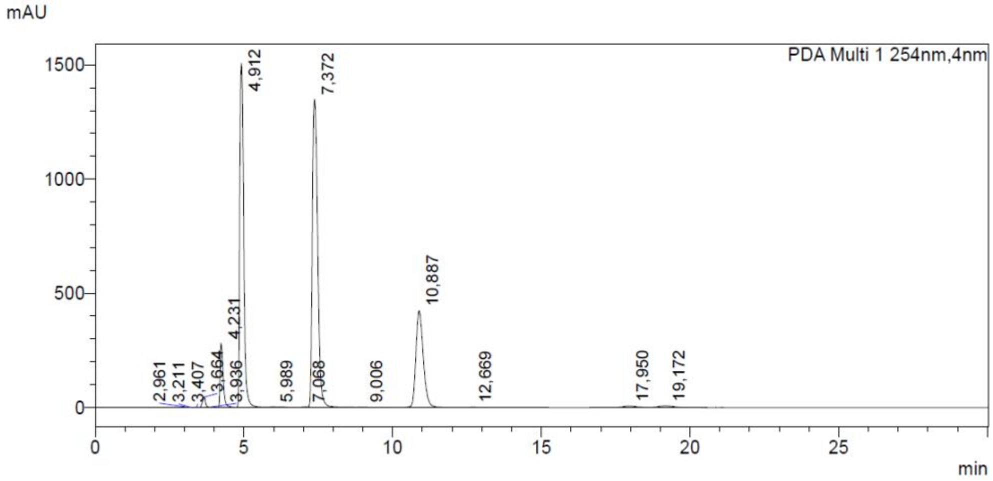


Sample from 6H5L_Top protein

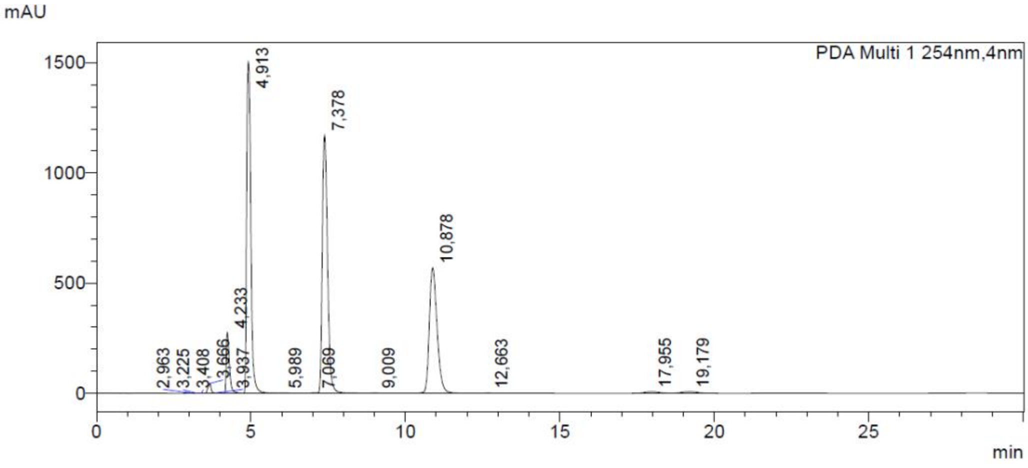

**Retro-aldol reaction**

Retention times were as follow: IS 4.9 min, **1** 7.5 min, **4** 11.1 min, (*R*/*S*)-**3** 18.4 min, (*S*/*R*)-**3** 19.6 min.

Sample from control reaction (Blank)

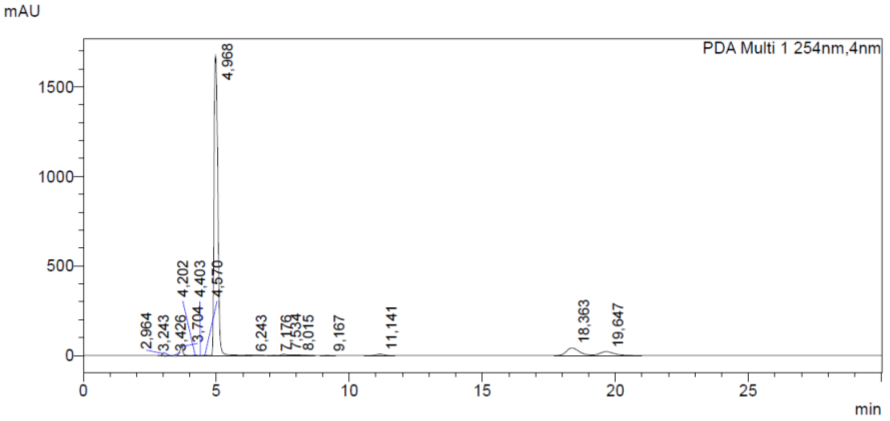


Sample from 6H5L protein

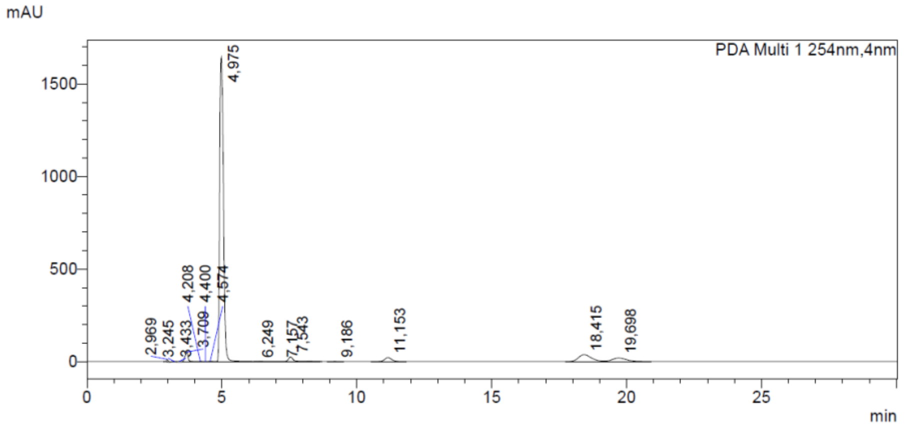


Sample from 6H5L_Top protein

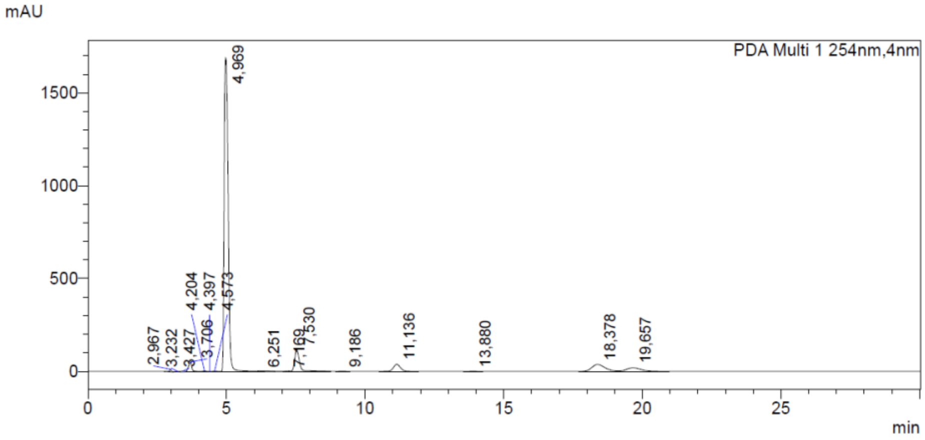

**Aldol reaction with reduced acetonitrile (1%) and acetone 2 (5%).**

Retention times were as follow: IS 4.9 min, **1** 7.4 min, **4** 10.7 min, (*R*/*S*)-**3** 18.5 min, (*S*/*R*)-**3** 20.0 min.

<u>Sample from control reaction (Blank)</u>

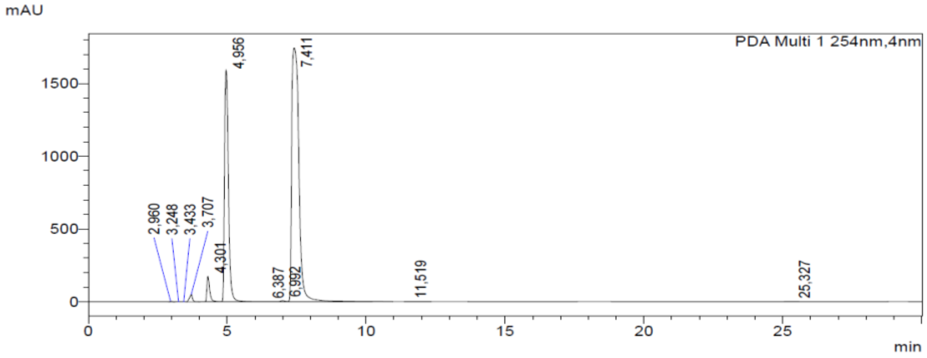


<u>Sample from 6H5L protein</u>

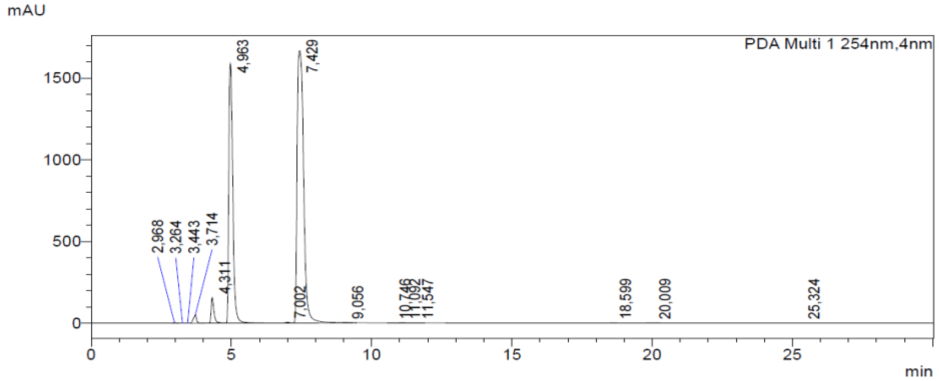


<u>Sample from 6H5L_Top protein</u>

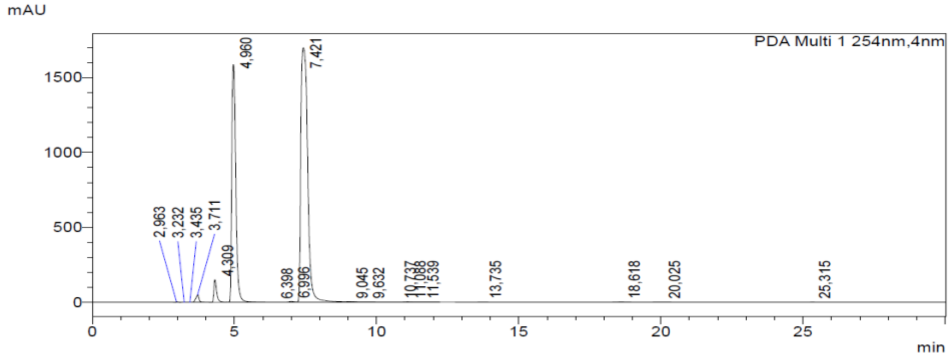

**Whole cells catalysis**

**Aldol reaction**

Retention times were as follows: IS, 4.9 min; 1, 15.4 min. The dehydrated product **4** and compound **3** co-eluted, appearing as a single peak at 19.7 min and could not be separated using this method.

Reference 6-methoxy-2-naphthaldehyde **1**

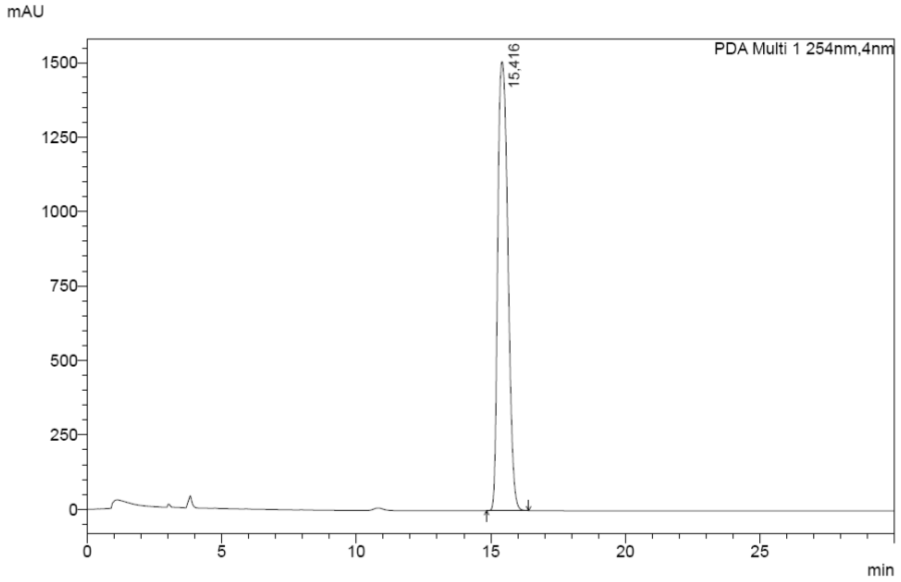


Sample from control reaction without cells

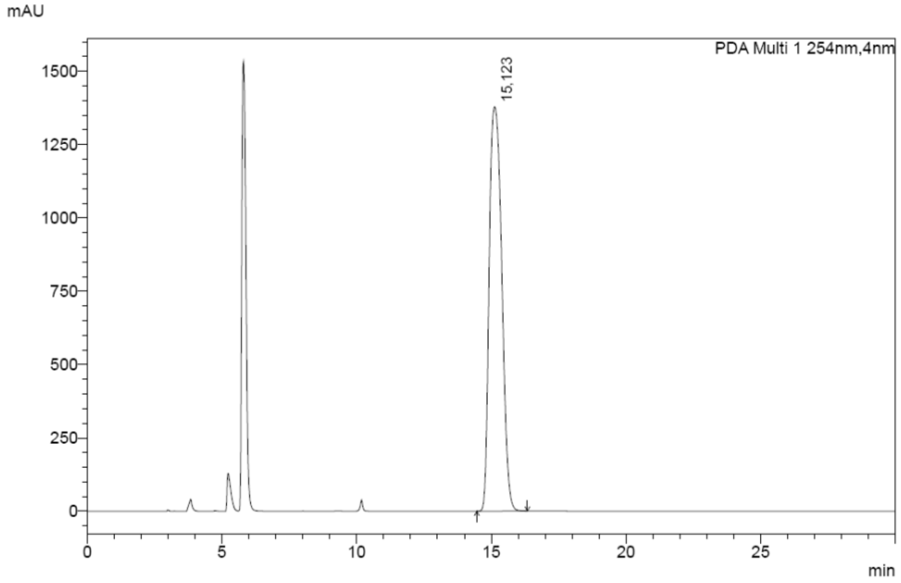

Sample from the aldol reaction with 6H5L_Cells

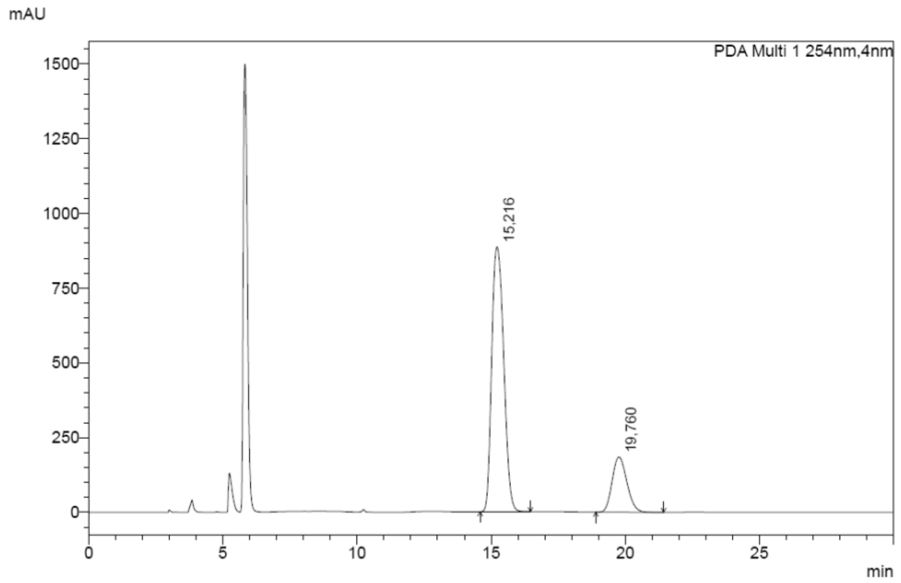


Sample from the aldol reaction with 6H5L_Top_Cells

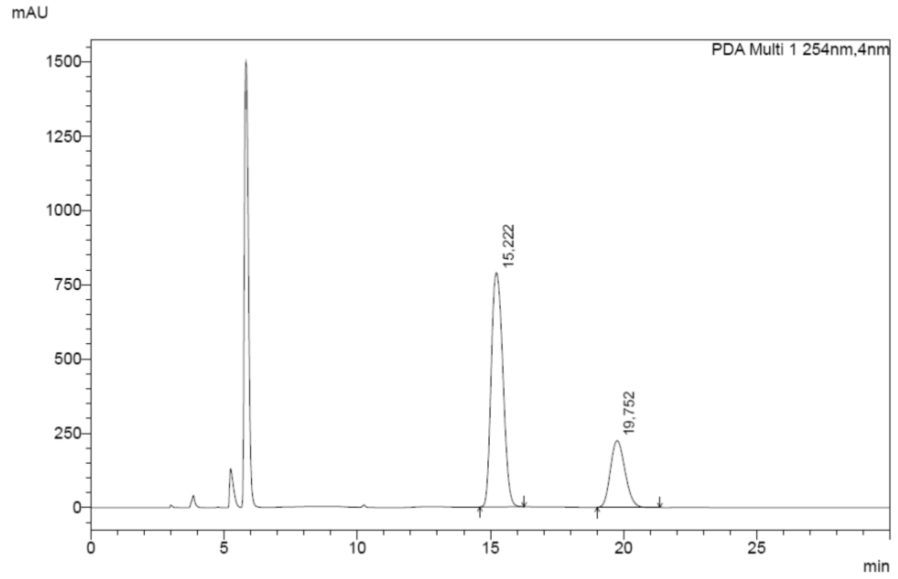

Sample from control reaction with empty cells (EnzFreeCells)

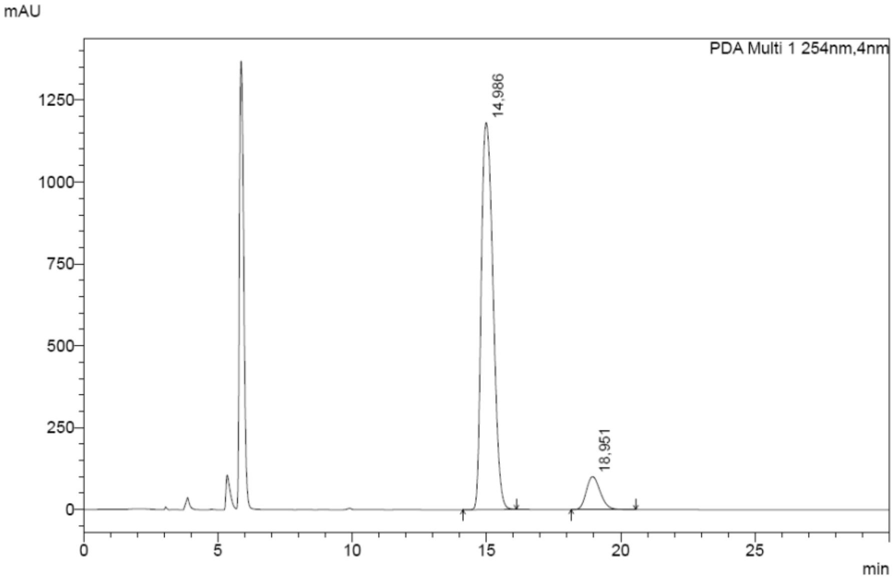


Sample from the aldol reaction with 6H5L Cells

Formation of *rac*-**3** along with the dehydrated product **4** was confirmed in the aldol reaction by changing the analysis on HPLC to the OD-H column, on which both compounds could be separated:

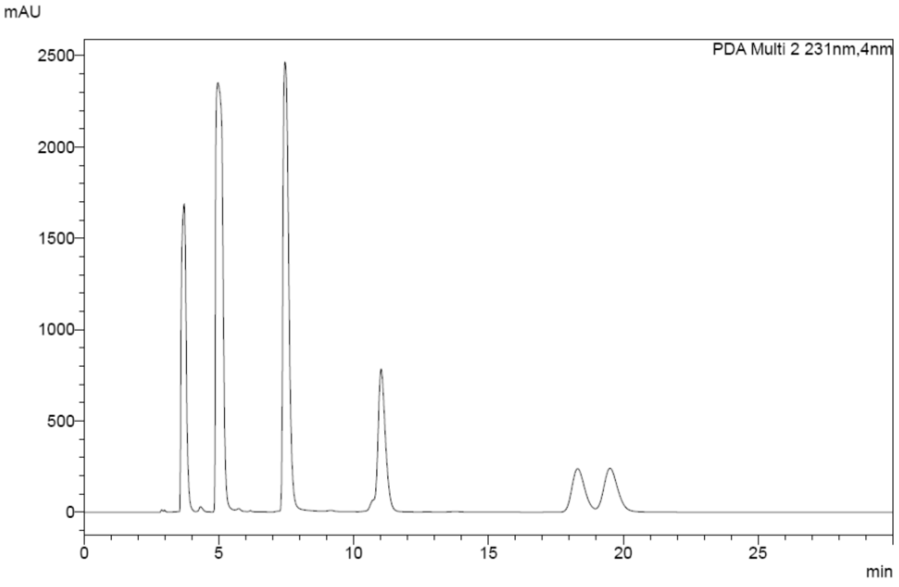

**Retro-aldol reaction:**

Retention times were as follow: IS 4.9 min, **1** 7.5 min, **4** 11.0 min, (*R*/*S*)-**3** 18.2 min, (*S*/*R*)-**3** 19.5 min.

<ins>Sample from control reaction without cells</ins>

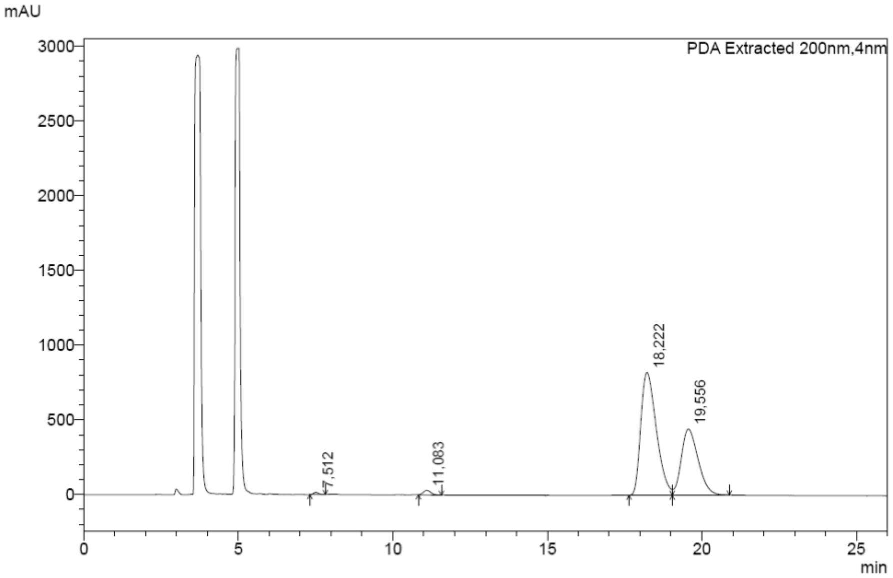


<ins>Sample from the retro-aldol reaction with 6H5L_Cells</ins>

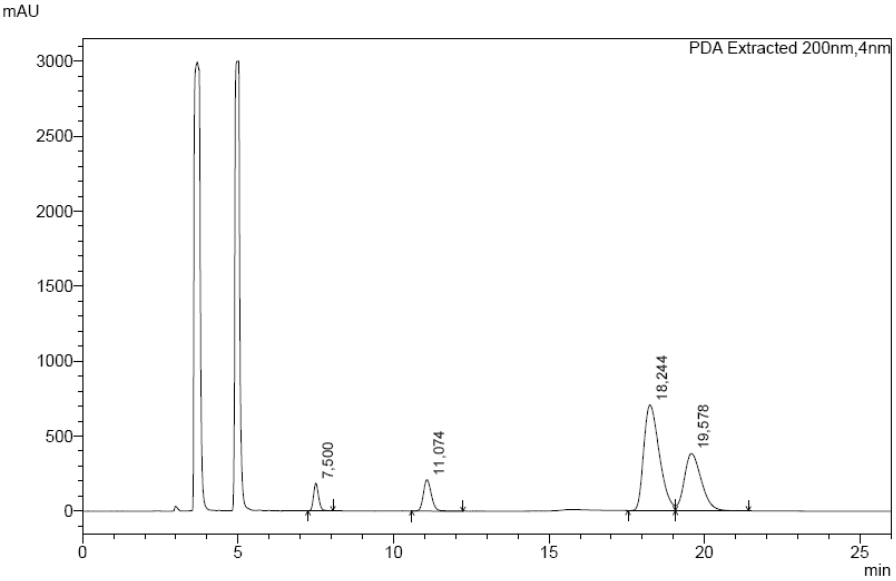

Sample from the retro-aldol reaction with 6H5L_Top_Cells

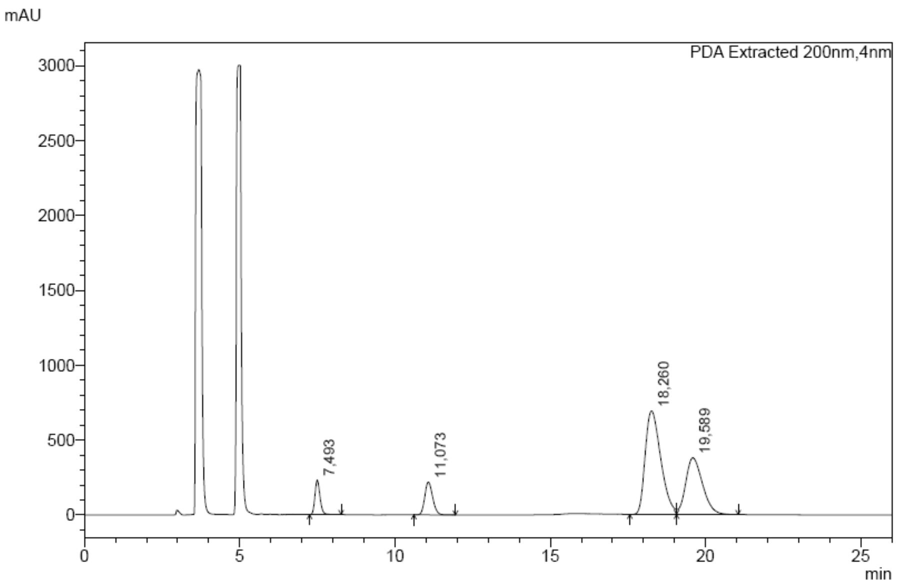


Sample from control reaction with empty cells (EnzFreeCells)

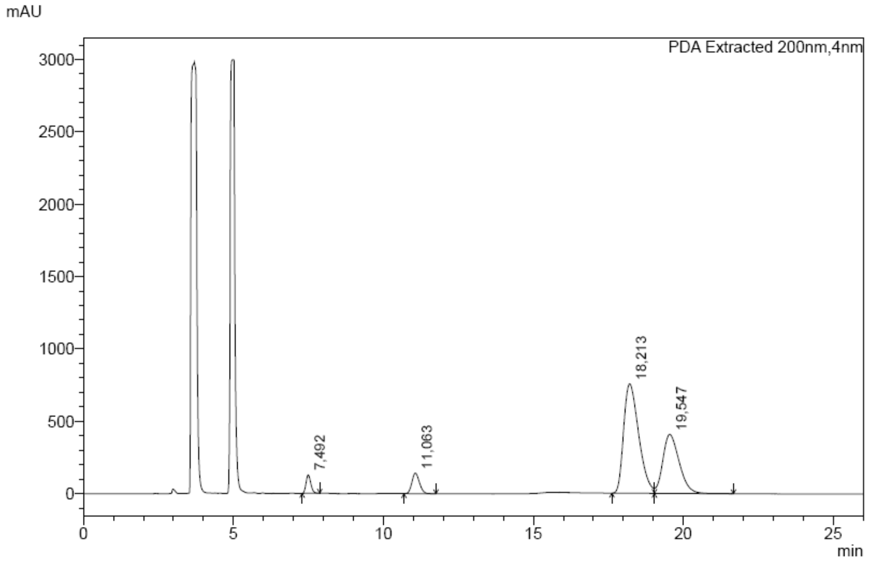

#### GC-MS analysis

The sample obtained from the aldol reaction with 6H5L was analyzed by GC-MS, which confirmed formation of the dehydrated product **4** (MW 226) with a retention time of 7.70 min (top: GC trace from GC-MS, bottom: mass fragmentation of peak eluting at 7.70 min).

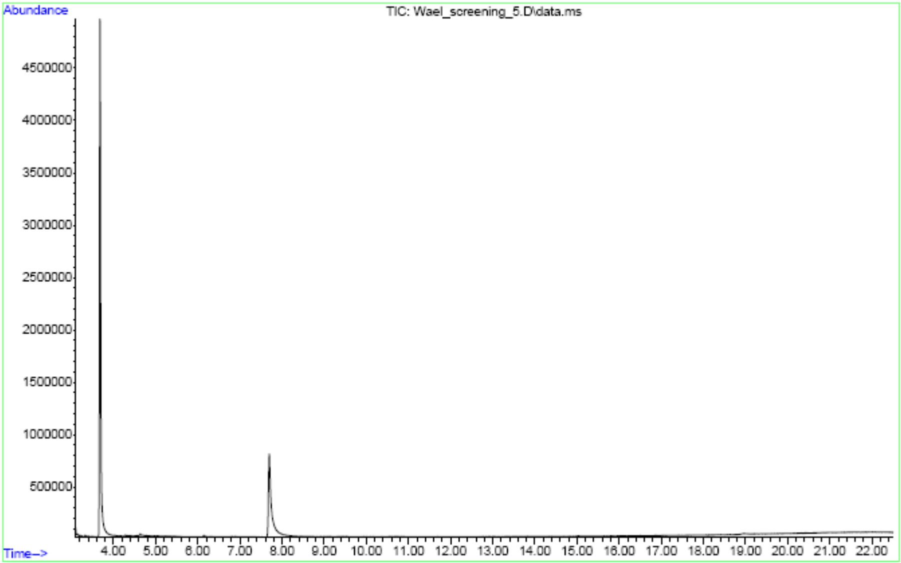


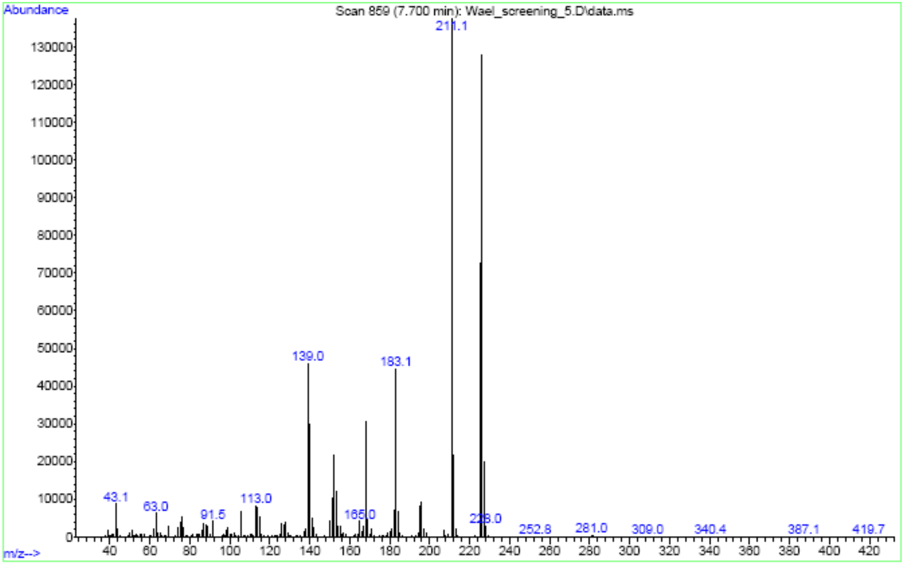

#### Stander curves

Standard curves of 6-methoxy-2-naphthaldehyde **1**, dehydrated product **4**, and (R/S)- and (S/R)-4-hydroxy-4-(6-methoxy-2-naphthalenyl)-2-butanone **3**.

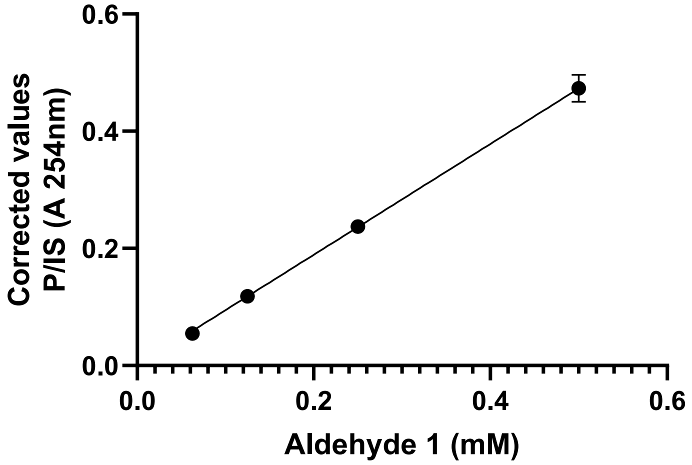


Y = 0.9462*X

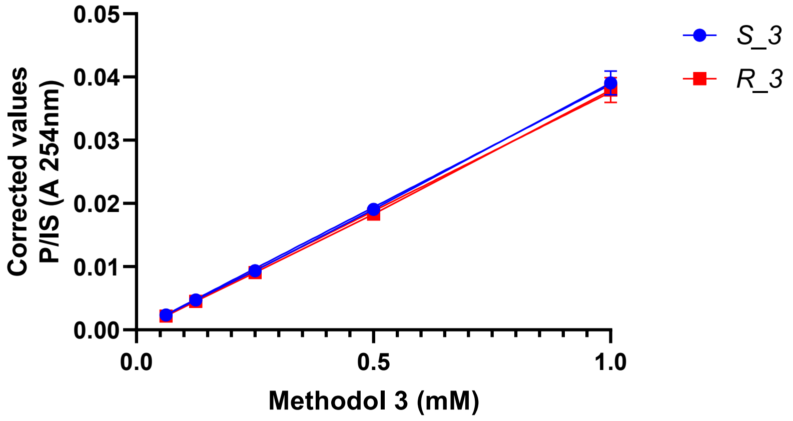


| *S*_3 | *R*_3 |
|---|---|
| Y = 0.03879*X | Y = 0.03759*X |

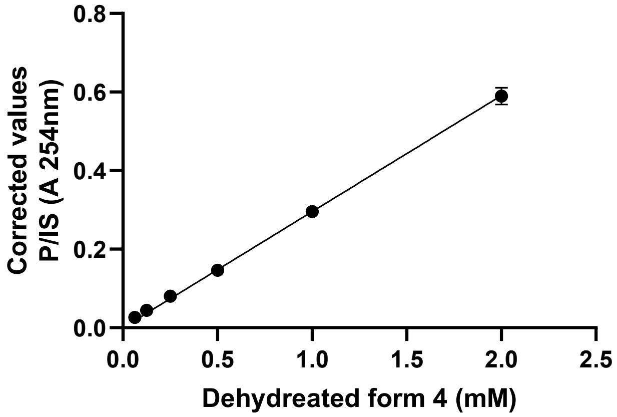


Y = 0.2955*X
